# Extended Plasmonic Nanostructures Templated by Tobacco Mosaic Virus Coat Protein

**DOI:** 10.1101/2024.12.26.630424

**Authors:** Ismael Abu-Baker, Alexander Al-Feghali, Elliot Zolfaghar, Gangamallaiah Velpula, Artur P. Biela, Steven De Feyter, Jonathan G. Heddle, Gonzalo Cosa, Amy Szuchmacher Blum

**Affiliations:** Department of Chemistry, McGill University, Montréal, Québec, Canada; Division of Molecular Imaging and Photonics, Department of Chemistry, KU Leuven, Celestijnenlaan 200F, 3001 Leuven, Belgium; Jagiellonian University, SOLARIS National Synchrotron Radiation Centre, Krakow, Poland; Jagiellonian University, Malopolska Centre of Biotechnology, Krakow, Poland; School of Biological and Biomedical Sciences, Durham University, UK

## Abstract

Optical and magnetic metamaterials possess interesting properties that cannot be achieved with conventional materials. However, there is currently no synthetic method offering both scalability and nanometer spatial precision. Biotemplating is a promising technique that has the potential to organize nanoscale components with high precision while being scalable and low-cost. Here we demonstrate a versatile template using hexahistidine-tagged tobacco mosaic virus coat protein. The protein self-assembles into disks which further assemble into extended nanostructures under mild conditions. Large sheets with either hexagonal or square packing and core-shell nanorods were formed, and gold nanoparticles were attached to the disks within each nanostructure to form assemblies of nanoparticle rings.

## Introduction

Nanomaterials show great promise in many advanced optoelectronic and magnetic applications, such as superlenses, data storage, nanoantennas, and catalysis.^1–4^ Functional nanoparticles with plasmonic, magnetic, or semiconductor properties have been synthesized with many morphologies and chemical compositions.^5–11^ However, control of particle composition and morphology alone is inadequate to achieve certain electromagnetic properties that arise from the coupling of electromagnetic fields between adjacent particles.^12–14^ Plasmonic and magnetic nanorings provide an example of intriguing properties that emerge from interparticle coupling, such as negative index of refraction and magnetic flux closure states.^15–17^ Rings of plasmonic particles exhibit coupled plasmonic modes that result in circular current and, consequently, an induced magnetic field. The electric and magnetic components of the refractive index can be tuned to achieve a negative index by controlling particle composition, spacing, and size.^18,19^ For a functional material, it is desirable to fabricate not just a single nanoring, but a surface of well-organized metal nanorings. Metasurfaces, consisting of planar periodic arrays of nanostructures, have received much attention due to the possibility of improved scalability and lower losses compared to complex 3D structures.^20^ Currently, most 2D metamaterials are made with block copolymer templates, lithography, or intrinsically layered materials, such as graphene and transition metal dichalcogenides.^21–24^ Common issues in existing systems include scalability, cost, and spatial precision.^25,26^

Many techniques have been developed to fabricate metasurfaces including block copolymer templates, lithography, biotemplates, and intrinsically layered materials, such as graphene and transition metal dichalcogenides.^21–24^ Common issues in existing systems include scalability, cost, and spatial precision.^25,26^ Traditional lithography, using extreme UV light or an electron beam to pattern a surface, has been expanded with high-throughput methods such as reflective electron beam lithography or interference lithography. However, these newer methods still require expensive equipment with complex optics, struggle to resolve features below ∼10 nm, and have limited material compatibility.^27–29^ Soft lithography methods, such as nanoimprint lithography, are cheap and high-throughput, but are still limited by the resolution of the initial mask/stamp, and challenges with certain pattern geometries.^30,31^ Block copolymer templates are affordable and scalable, but limited by polymer synthesis, complex phase behaviour, and difficulties in creating complex patterns.^32–34^ Liquid crystals have also been explored for bottom-up self-assembly of nanomaterials. Applications of liquid crystals are often limited by the requirement for high concentrations of pure, monodisperse components, and difficulties in templating certain shapes, including rings, although some examples of large rings have been reported.^35–37^

Biotemplating has potential as a method to organize nanoscale components with high precision. Many biological nanomaterials have been reported to template metal nanostructures. These include DNA origami, peptides, virus-like particles, and other proteins.^38–41^ Depending on the template, advantages include customizability, scalability, sustainability, low cost, and high spatial precision. DNA origami has received the most attention, and while it does provide very high precision and reproducibility, scale-up is limited by current DNA synthesis and sequencing technologies, and precise functionalization with proteins or metals remains challenging.^42–45^ Tobacco mosaic virus coat protein (TMVP) is particularly appealing for the formation of metal nanorings. At near neutral pH, TMVP self-assembles into disk-shaped particles with a diameter of 18 nm.^46^ The circular nature of the disks makes them an ideal nanoring template. The protein can be produced in large quantities in plants, yeast, or *E. coli*.^47,48^ Self-assembly from monomer to disk occurs under mild conditions, and disks self-assemble into helical rods below pH 6.5. A number of mutants have been created to modify the assembly behaviour and chemical functionality of the disks/rods.^49–52^ Because the disk structure is well characterized, genetic engineering can be used to add functionality to specific locations on the disk, allowing precise attachment and organization of nanoparticles and adaption of the template to different nanoparticle compositions.^53^

Previous work demonstrated several methods to couple metal nanoparticles to the outer edge of TMVP disks to form nanorings. These commonly involve cysteine or lysine mutations at the outer radius.^54,55^ A hexahistidine-tagged TMVP mutant was shown to self-assemble into large hexagonally packed sheets of disks.^56^ Zhang *et al*. took this approach a step further and demonstrated large hexagonally packed sheets of TMVP disks with gold nanoparticles (AuNPs) attached to form sheets of metal nanorings.^54^ This assembly was created with a tetrahistidine-tagged mutant in which Cu^2+^ chelation by the His-tag was used to assemble the protein sheets, and AuNPs were subsequently bound to the His-tag. Ag_2_S and CdSe@ZnS nanoparticles were also able to bind to the protein template. Here we report the formation of two novel extended nanostructures (square packed sheets and core-shell nanorods) from TMVP with a C-terminal hexahistidine tag (6H-TMVP). These, along with extended hexagonal assemblies, were used to organize AuNPs into nanorings without metal chelation, generating nanostructures with extended plasmon resonances via self-assembly on scales previously achieved only via lithography.

## Results and Discussion

### Formation of regularly shaped, hexagonal sheets

6H-TMVP was found to self-assemble into a diverse array of nanostructures, many of which have not been previously reported. The protein has a strong preference to form 3- and 4-layer disks in addition to the canonical bilayer disk (Fig. 1). Although 3- and 4-layer disks have been reported previously, 6H-TMVP is noteworthy for forming these disks as the primary disk species in low ionic strength and from pH 5.0-10.0. 6H-TMVP disks further self-assemble into rods or extended nanostructures depending on pH, ionic strength, protein concentration, and solution additives.

**Figure 1.**
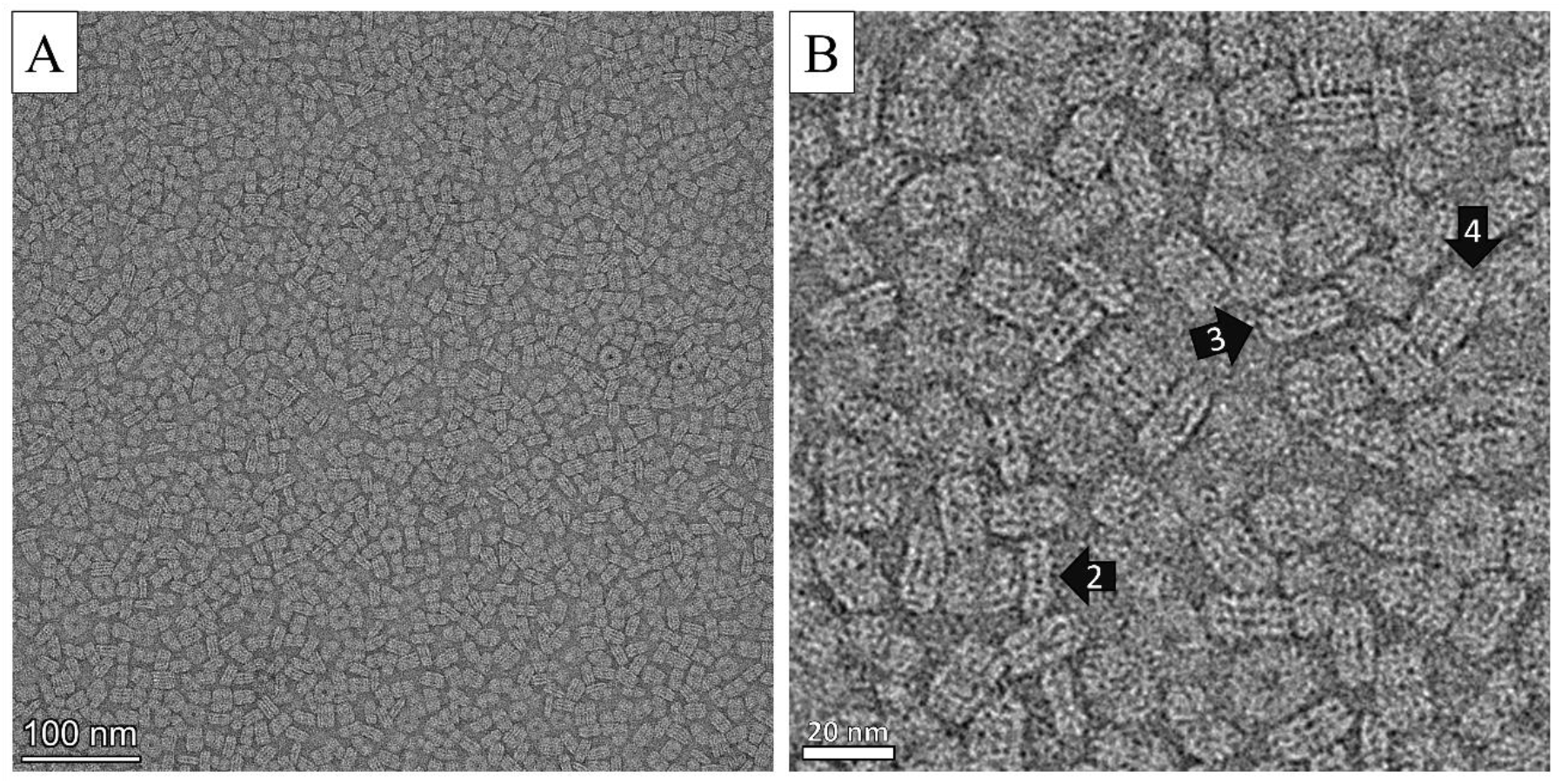
(A) Representative TEM image of 6H-TMVP disks showing primarily 2-, 3-, and 4-layer disks. (B) Zoom-in with arrows indicating 2-, 3-, and 4-layer disks.

Hexagonally packed sheets of disks were observed from pH 5.0-10.0. Phosphate buffer in the pH range 6.0-6.5 led to the fastest formation of large sheets. These sheets could be single or multilayered and measured up to several microns across (Fig. 2A, B). At potassium phosphate concentrations above 150 mM, stacked disks dominate and form low quality hexagonally packed clusters. Rods become the dominant species below pH 6.0, and sheet formation at alkaline pH is extremely slow. Initial experiments at pH 6.8 in 100 mM potassium phosphate buffer showed significant kinetic trapping of the disks during sheet formation resulting in poorly formed sheets with irregular shapes and many defects (Fig. 2C). Small hexagonally packed sheets were observed in cryogenic electron microscopy (cryo-EM) screening of samples.

**Figure 2.**
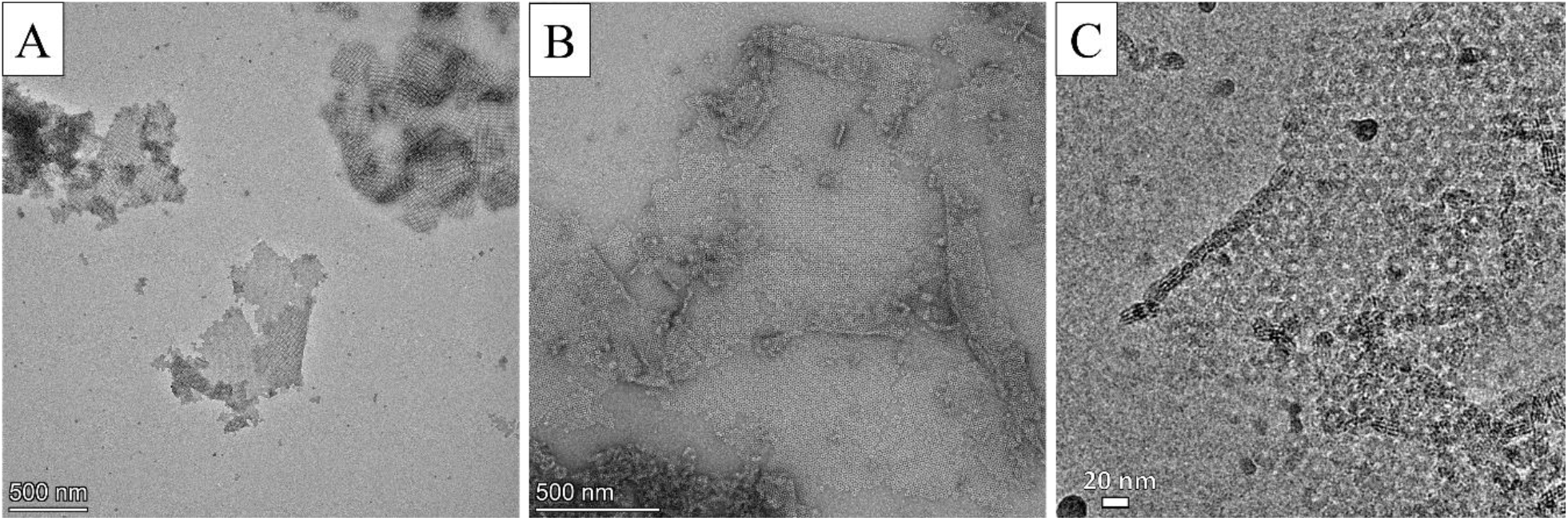
TEM images of hexagonal 6H-TMVP sheets. (A) Irregularly shaped sheets due to kinetic trapping. (B) Higher magnification image of hexagonal sheets. (C) Sheets observed in cryo-EM.

Inclusion of a high concentration of imidazole during assembly was found to yield large, regularly shaped, hexagonal sheets with few defects (Fig. 3A, B). This effect appears to depend on the ratio of imidazole to protein. As imidazole mimics the histidine side chain, imidazole-His interactions likely compete with the His-His interactions involved in sheet formation, making disks that bind to a sheet in a non-close-packed position less favourable. At lower ratios of imidazole to protein, high aspect ratio bundles formed from rods/stacked disks (Fig. S1). Similarly, the presence of 5.0 mol% ethanol during dialysis yielded large sheets with few defects (Fig. 3C, D). Lower concentrations of ethanol were increasingly less effective, but higher concentrations caused large aggregates of randomly oriented disks. Overall, 5.0 mol% ethanol was found to be the simpler and more cost-effective way to improve hexagonal sheet formation. Unlike imidazole, no alternative assembly pathways were observed with ethanol-mediated assembly, regardless of protein concentration.

**Figure 3.**
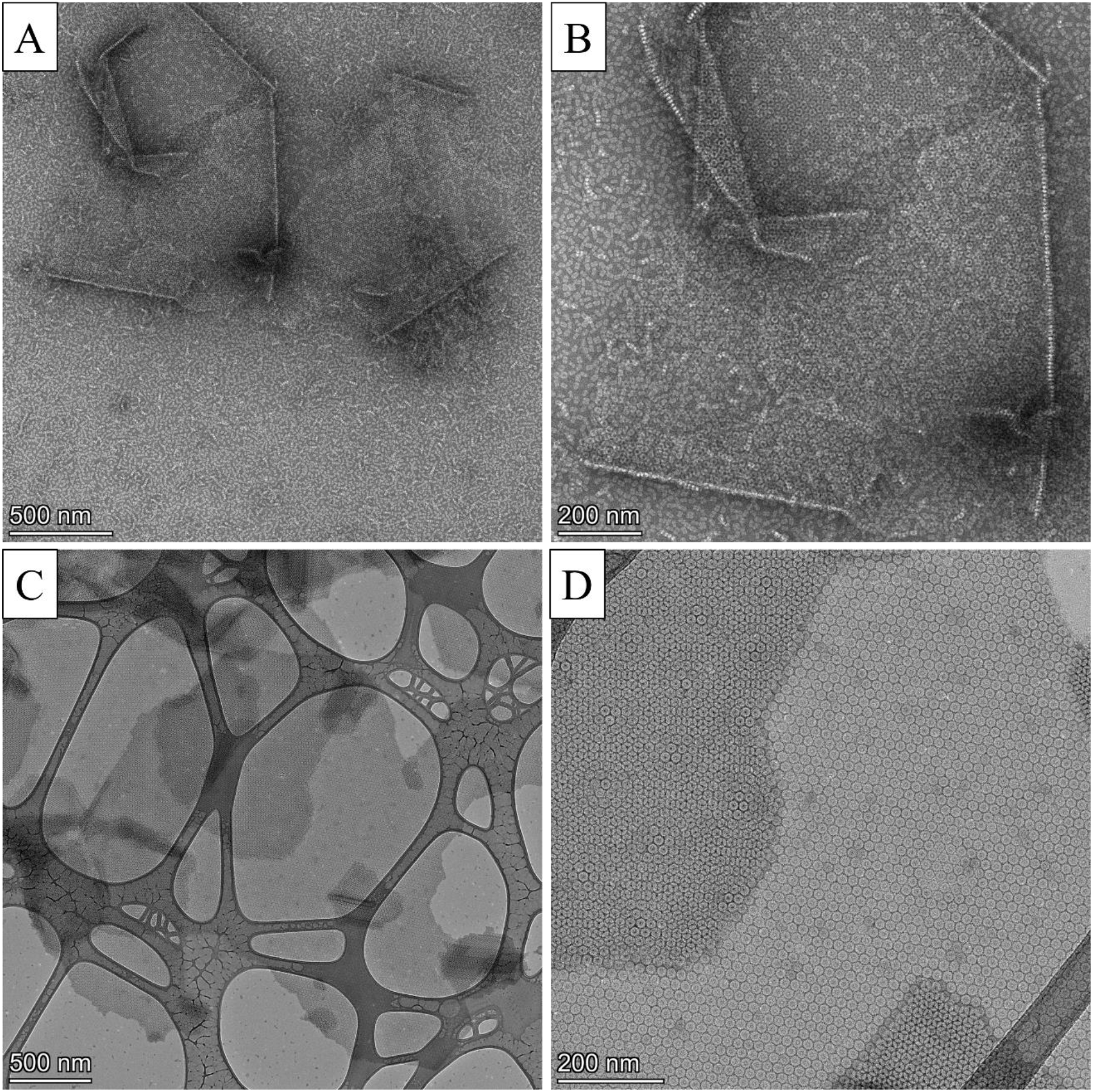
TEM images of nanostructures formed in the presence of imidazole or 5.0 mol% ethanol. (A) large hexagonal sheets formed with imidazole. (B) Higher magnification image of A. (C) Low magnification image of sheets formed in ethanol. (D) High magnification image of C showing 1- and 2-layer regions.

### Templating AuNPs on square and hexagonal sheets and nucleating new sheets

AuNPs capped with Bis(p-sulfonatophenyl)phenylphosphine (BSPP) were attached to hexagonal sheets by mixing an excess of AuNPs with the sheet solution and incubating overnight. Alternatively, AuNPs could be mixed with unassembled disks under the same solution conditions to trigger assembly of hexagonally packed sheets. In this case, the strong interaction between gold and histidine causes rapid formation of low-quality sheets. Incubating AuNPs with 6H-TMVP at pH 6.0-6.3 in 50-100 mM potassium phosphate at 4 ^°^C resulted in formation of sheets with square packing (Fig. 4A, B). Micron-sized multilayer sheets formed in solution within 15 minutes of mixing protein and AuNPs at 4 ^°^C (Fig. S2). Formation of square lattices is more favourable at protein concentrations near 2.5 mg/mL, but hexagonal sheets become dominant at low protein concentrations. The protein must be maintained at low temperature to avoid hexagonal sheet formation which occurs quickly at room temperature. Most square sheets were in the range of 1-5 µm but sheets up to 10x10 µm were observed in TEM (Fig. 4A, B). Multilayer square sheets tend to exfoliate into single layer sheets under the capillary force during TEM grid preparation (Fig 4C). AFM of a dried multilayer sheet showed a smooth surface on the top layer with a step height of approximately 4 nm between each layer (Fig 4D). Square sheets were not formed in control experiments in which the AuNP solution was replaced with an equal volume of BSPP solution (Fig. S2C).

**Figure 4.**
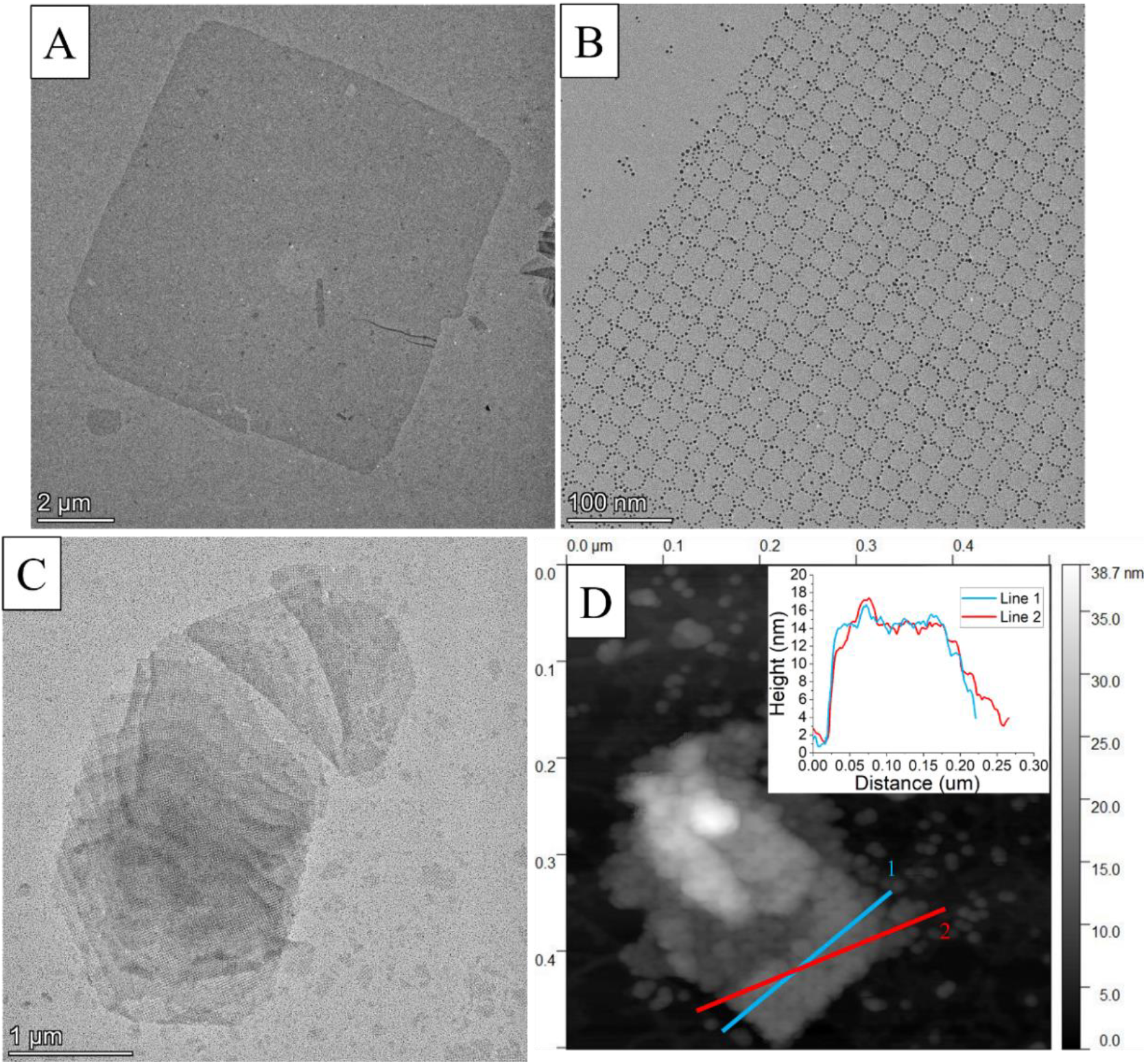
Images of square sheets. (A) TEM image of a very large single-layer square sheet. (B) High mag image of square sheet where AuNP arrangement can be observed. (C) TEM image showing exfoliation of a multilayer square sheet. (D) AFM image of a square sheet with height profile inset.

Additionally, when protein and AuNPs were mixed directly on a TEM grid, only hexagonal sheets were observed, indicating that the square sheets form in solution and are not nucleated by the grid surface. Sheets frequently appear folded or crumpled in TEM and AFM images, further supporting the idea that these sheets form in solution (Fig. S3). Protein-only square sheets were observed on rare occasions, but no reliable protocol to form them could be determined. 6H-TMVP incubated under standard square sheet conditions did not assemble into any extended structures after two months at 4 ^o^C without the addition of AuNPs. However, 6H-2R-TMVP, a mutant that does not form helical rods due to K53R and K68R mutations, showed a stronger preference to adopt square packing in the absence of AuNPs, especially large body-centered tetragonal (BCT) crystals in which each layer has square packing (Fig. S4).^51^

Both 3 and 5 nm AuNPs could be templated with this method. Sheet quality appears to decrease as nanoparticle size increases, with 8 nm AuNPs forming low quality sheets with many defects (Fig. S5). Lattice constants, measured by TEM, were the same for all the observed sheets with or without AuNPs attached (Table. S1). This indicates that the hexagonal and square sheets are formed by similar interactions and that AuNPs are not in the plane of the disks, even when AuNPs are used to trigger sheet formation. If AuNPs were bridging between disks without protein-protein interactions between neighboring disks, the lattice constants would be larger with AuNPs bound. Interestingly, AuNPs appear to only attach to a single face of the sheet in both hexagonal and square lattices. This suggests that either 6H-TMVP disks lack dihedral symmetry under the conditions used for assembly or that AuNP binding modifies the structure of the bound face in a highly cooperative process. Based on preliminary cryo-EM results, approximately 40% of the disks are 3-layer and 50% are 4-layer at pH 6.0 (Fig. S6). Both of these disks appear similar to the wild-type (WT-TMVP) 4-layer aggregate from the literature.^53^ While the type of disk in a typical sheet could not be determined, the tendency for nanoparticles to only bind to one face of the sheets could indicate that the sheets are primarily composed of 3-layer disks, which do not have *C*_*2*_ symmetry. A full cryo-EM comparison of the wild-type and multiple His-tagged TMVP mutant disks is underway to better understand how these mutations affect the disk structure.

TEM images of square and hexagonal sheets shortly after mixing with AuNPs revealed very different assembly behaviour for the two structures. In both cases, AuNPs show a strong preference to first bind at the vertex of multiple disks and the outer edges of sheets. The outer edges of sheets have the most mobile and least sterically hindered histidine residues, making AuNP binding favourable. The lattice vertices offer the highest number of histidine binding partners for the AuNPs, making them the most favourable binding locations within the sheets. In the case of hexagonal, this leads to initial binding of six AuNPs with the rest of the ring quickly filled in afterwards (Fig. S7A). Shortly after initiating assembly, square sheets with AuNPs only along the outer edge and sheets with AuNPs at each vertex where 4 disks meet were both observed (Fig. S7B, C). It is unclear if either or both of these partially assembled structures represent the nucleating structure. This difference in initial AuNP binding may contribute to the tendency for square sheets to have larger spacing between AuNPs, and therefore less plasmonic coupling between particles. In addition to single-layer sheets with incomplete nanoparticle binding, multilayer stacks of large square sheets that do not appear to have any AuNPs specifically bound were observed (Fig. S7D). Given that square sheets do not form under the same conditions without the addition of AuNPs, these protein-only stacked sheets are assumed to be nucleated by sheets with bound AuNPs. Once AuNPs nucleate one square sheet, that sheet can nucleate the growth of square sheets without any AuNPs, and those protein-only sheets can continue to nucleate more sheets, creating multilayer stacks without bound AuNPs. Within 48 hours, the multilayer protein-only sheets fully bind rings of AuNPs. The multilayer sheets with AuNPs attached exfoliate during TEM grid preparation, while the protein-only sheets do not. This indicates that AuNPs weaken the interlayer interactions. pH measurements confirmed that the solution pH did not change after adding AuNPs.

### Formation of core-shell nanorods and their templating of AuNPs

When assembled at pH 6.0 in 50 mM potassium phosphate and 3.5 mol% ethanol, 6H-TMVP formed core-shell nanorods (CSNRs) which consist of a hexagonal sheet of disks wrapped around a helical TMVP rod (Fig. 5). The structure is metastable and extremely sensitive to assembly conditions, which hindered many characterization methods such as cryo-EM. CSNRs were only observed at pH 5.8-6.0, 75-150 mM ionic strength, and 0.75-1.5 mg/mL protein concentration. CSNRs assembled in MES and bis-tris buffers with added NaCl, but assembly was most consistent in phosphate buffer. Similarly, CSNRs could be formed without ethanol, but the assembly was found to be much more consistent in the presence of 3.5 mol% ethanol. We have previously shown that a similar concentration of ethanol prevents the transition from disk to helical rod in wild-type TMVP.^57^ Ethanol may similarly affect 6H-TMVP self-assembly to help maintain appropriate populations of both disks and rods. The His-tag increases the propensity to form helical rods, so this concentration of ethanol is not able to completely prevent rod formation as in WT-TMVP.

**Figure 5.**
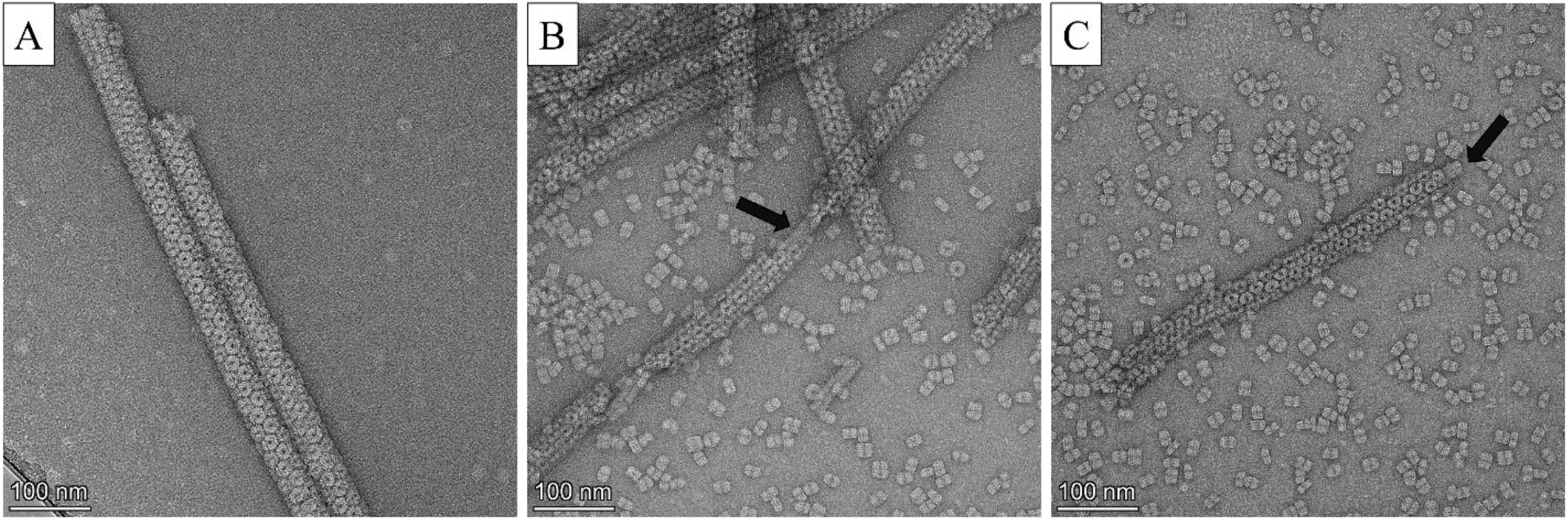
TEM images of core-shell nanorods. Arrows point to helical rods that appear to form the core of the CSNR.

TEM images frequently show helical rods protruding from the ends of CSNRs as well as staining that matches the presence of a helical rod inside the CSNRs (Fig. 5B, C). While most CSNRs appear to have a single helical rod in the center, there is a small population with larger diameters that seem to have a small bundle of rods in the center. Some structures have also been found to be completely lacking a helical rod, creating a hollow nanotube. These nanotubes do not have a large enough inner diameter to accommodate a helical rod. Negative stain tomography showed that the nanotubes collapse upon drying (Movie S1).

While it is unclear if these narrower nanotubes result from CSNRs that grow beyond the length of the helical rod or if they nucleate from a sheet of disks without rod involvement, we suggest the former mechanism because no CSNRs or nanotubes were observed under the same conditions using 6H-2R-TMVP, which has mutations that prevent helical rod formation. The hexagonal lattice was observed to have two possible orientations relative to the tube axis. Using the analogous carbon nanotube notation, CSNRs/nanotubes can have armchair or zigzag lattices (Fig. S8). CSNRs with a single helical rod in the center always had the zigzag structure, while assemblies with zero or multiple rods could adopt zigzag or armchair structures. Based on TEM images, the disks in core-shell nanorods appear to primarily be 4-layers, although layers could only be counted for disks on their edge, so the number of layers in most disks in any given CSNR could not be determined. As with the TMVP sheets, AuNPs can be attached to CSNRs/nanotubes to create gold nanorings in a tubular hexagonal lattice (Fig 6).

**Figure 6.**
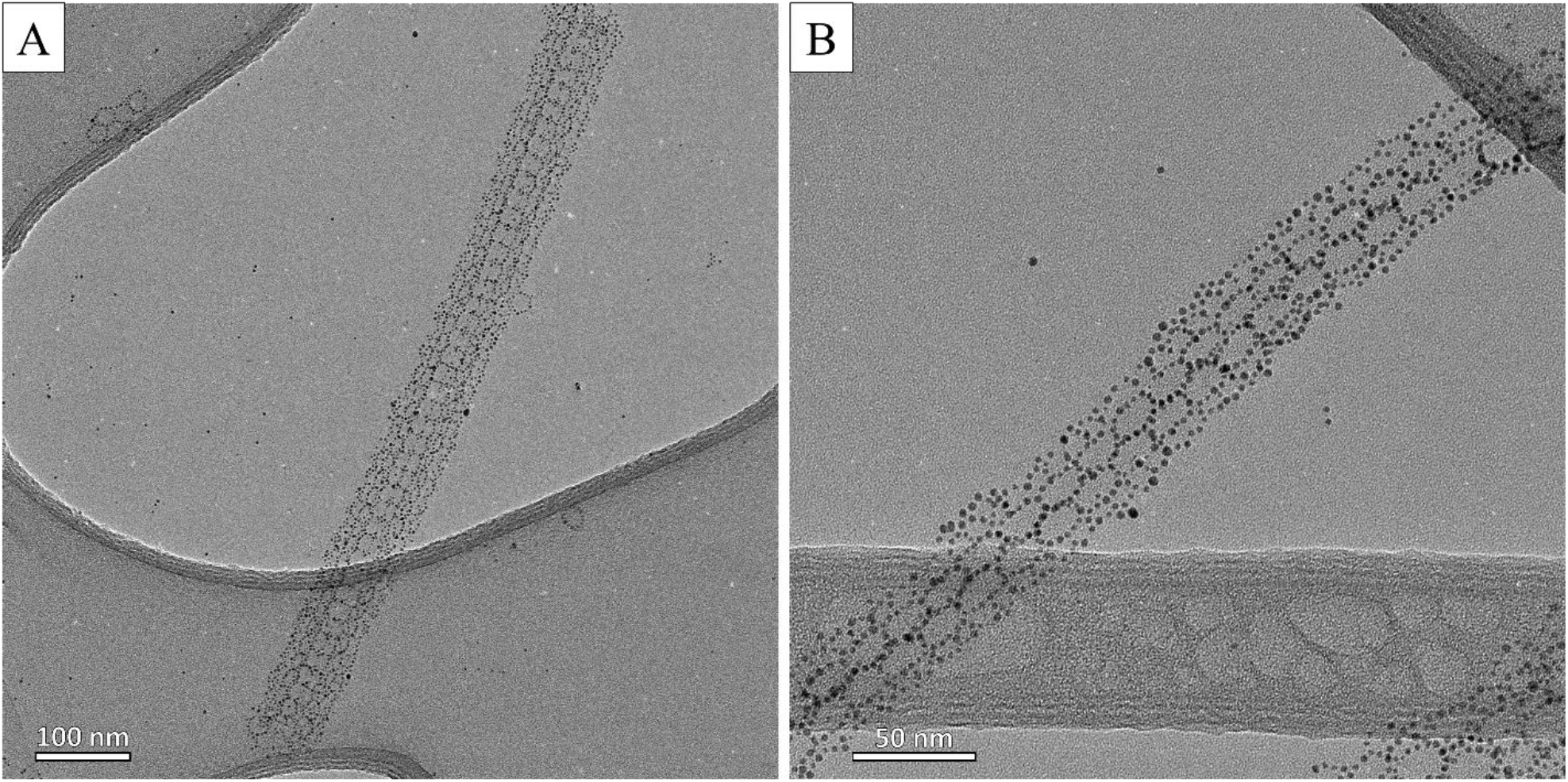
TEM images of 6H-TMVP nanotubes/CSNRs with AuNPs attached.

Histidine has primarily been studied as a transition metal chelator, and many examples of materials self-assembled through metal-His interactions exist.^54,58–60^ To investigate effects from possible metal contamination, assembly was attempted in various concentrations of ethylenediaminetetraacetic acid (EDTA). No changes were observed in 20 mM EDTA (data not shown). Assembly of square and hexagonal sheets in high concentrations of EDTA strongly resembled the results of assembly in high salt conditions (Fig. S9-11). As EDTA is an acid, neutralizing high concentrations of EDTA led to a significant increase in the ionic strength. Considering that similar effects were observed from high concentrations of EDTA and NaCl, it seems likely that changes in assembly were primarily caused by the increased ionic strength rather than removal of contaminating metal ions. Comparison to 4H-TMVP, which has a shorter histidine tag, shows that the number of histidines is crucial to sheet assembly. Unlike 6H-TMVP, a 4H-TMVP mutant was unable to form extended assemblies, even in the absence of EDTA. This mutant was previously shown to form large hexagonally packed sheets through Cu^2+^ chelation.^54^ The self-assembly of 6H-TMVP but not 4H-TMVP suggests that the assemblies reported here rely on a large number of histidine-histidine interactions rather than metal coordination.

While all assemblies could be formed in non-phosphate buffers, such as bis-tris, MES, and MOPS, assembly was found to be significantly more favourable and consistent in potassium phosphate buffer, even when controlling for ionic strength. One possible explanation for this is that assembly occurs partially through His-phosphate interactions. This may be through salt bridges or hydrogen bonding. Hexagonal sheets can form at pH values ranging from 5.0-10.0, so assembly can occur with histidine in any protonation state. In the case of CSNRs, histidine residues on the helical rod likely interact with residues on the face of the disk, which includes arginine and carboxylic acids, further adding to the possible bridging interactions with phosphate. Supplementing bis-tris buffer with sodium sulfate did not replicate the effect of phosphate on assembly, instead causing poorly ordered aggregates of rods and hexagonal sheets to form (Fig. S12). Phosphate has a pKa of 7.20, which would allow it to participate in more dynamic interactions, while sulfate does not have a pKa within the range of this study. His-His interactions have been shown to stabilize some proteins, but few studies have demonstrated the significance of His-His interactions in large-scale quaternary protein assemblies.^61–63^ It appears 6H-TMVP self-assembles through many weak histidine interactions, leading to rapid self-assembly of extended nanostructures with few defects and tunable structures based on assembly conditions.

### Hyperspectral darkfield of templated of AuNPs

Due to the large size and poor colloidal stability of these self-assembled structures, typical solution-based UV-vis absorption spectra do not capture the behaviour of individual nanostructures. To both characterize these large structures and compare signals arising from distinct and separate assemblies, we collected hyperspectral darkfield microscope images of both square and hexagonally packed AuNP-6H-TMVP assemblies deposited on carbon-supported pinpointer TEM grids (Fig. 7). 3 nm gold nanoparticles have a very small scattering cross-section, and thus are expected to be invisible in dark field scattering measurements. Control experiments performed on 3 nm nanoparticles alone show no signal. In contrast, both structures show a bright edge with a dim interior (Fig. S13). This is likely due to symmetry breaking at the edges, resulting in strong responses at resonance.^64^

**Figure 7.**
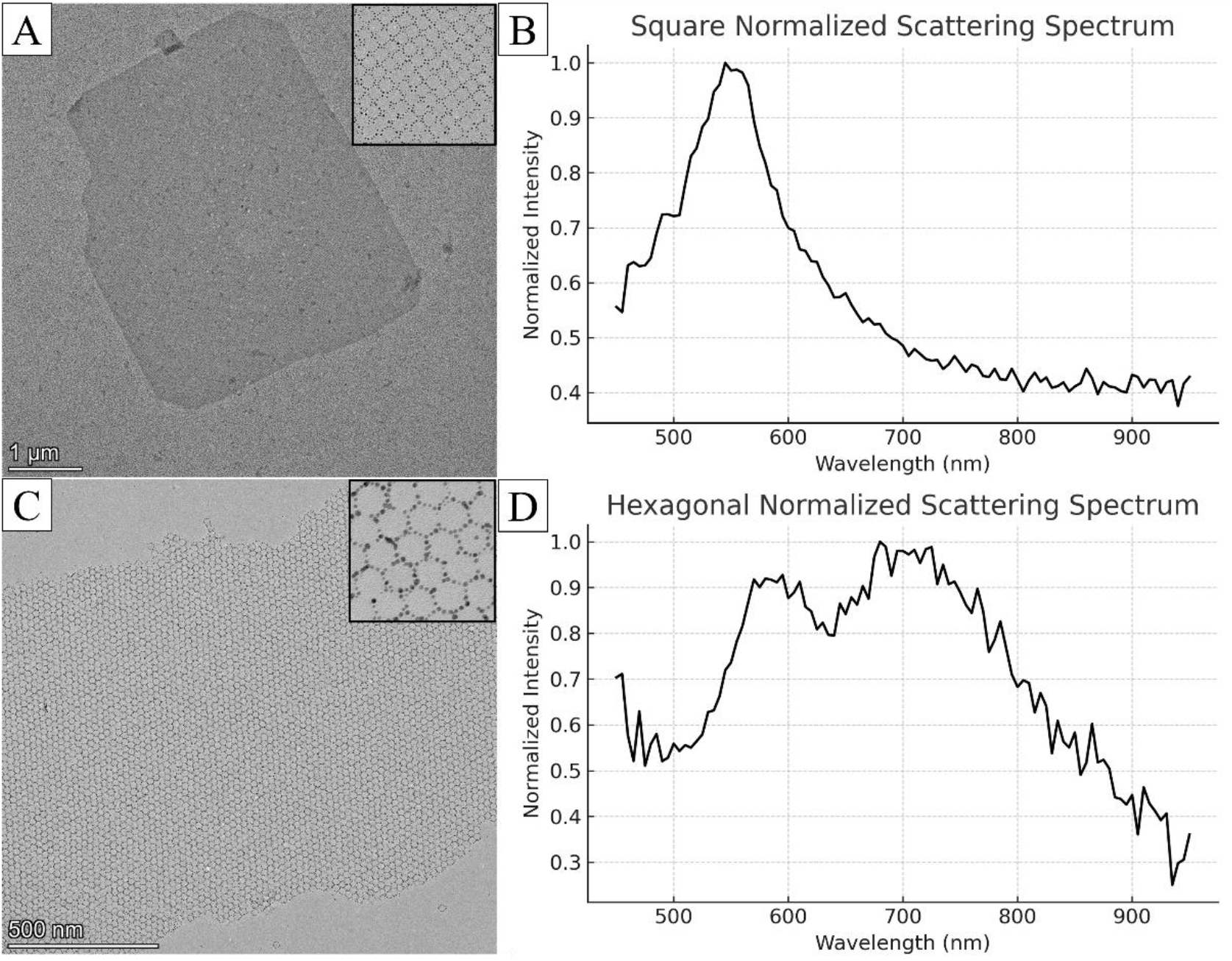
(A) TEM image of square sheets that were measured in darkfield microscopy. (B) Corresponding darkfield scattering spectrum of square sheets. (C) TEM image of hexagonal sheets that were measured in darkfield microscopy. (D) Corresponding darkfield scattering spectrum of hexagonal sheets.

The square-packed assemblies of 3 nm Au NPs exhibit a strong peak at 520 nm, which is red-shifted slightly compared to the resonance of individual 3 nm AuNPs at ∼514 nm (Fig. S14). In contrast, the hexagonally packed assemblies consistently exhibit a redshifted response, indicating a different type of coupling between nanoparticles. The lower nanoparticle density of the square assemblies compared to hexagonal assemblies might explain this difference in optical behaviour, where the larger size of square vs hexagonal may play no role in the boundary conditions leading to the observed spectra. For 3 nm nanoparticles, small differences in interparticle distance result in large changes in optical properties.^65^ While these spectra correlate with individual sheets, differences in interparticle distance, packing geometry, and sheet-to-sheet interactions preclude a detailed analysis. In addition, the sizes of many structures correspond to the diffraction limit, further complicating comparison of the edges with the center. High magnification images and some particle/sheet statistics are presented in Fig. S15, S16 and Table S1. Further experiments and simulations are currently underway to better understand the optical response of the sheets.

## Conclusions

In this study we demonstrated a versatile method to template extended metal nanostructures using 6H-TMVP disks. Histidine-based assembly without metal chelation allows the system to adopt multiple structures depending on solution conditions. This is the first report of square sheets or CSNRs from TMVP. Although core-shell nanorods have been reported for polymer, protein, and DNA systems, to our knowledge, a CSNR in which the core and shell are composed of the same protein assembled into different types of virus-like particles has never been reported before.^66–68^ Nanostructures appear to form in solution, so deposition on any desired surface should be possible. With further study, assembly of sheets directly on surfaces is likely possible as well.^69^ Gold nanoparticles were bound to the protein template through interaction with the His-tag. Histidine-based attachment of protein to gold, silver, and ZnS nanoparticles has previously been demonstrated.^70–73^ A thorough investigation of histidine interactions with a wider range of nanoparticle compositions would be beneficial to the biotemplating field. Square sheets tend to show lower AuNP packing density in each ring, which reduces plasmonic coupling across the structure. The difference in packing is likely due to initial binding of four AuNPs on each disk of the square lattice rather than the six AuNPs that bind at the vertices of hexagonal sheets. Adjustments to nanoparticle size and ligands as well as the protein template may improve the plasmonic coupling in the sheets in the future.

The protein template can also be engineered to display other residues, such as cysteine or lysine, to allow binding of nanoparticles or biomaterials that do not interact well with histidine. This would enable viral templates for use with materials that are not suitable for harsher methods like lithography. For real-world application of these templated nanostructures, it would be advantageous to create mutants that are optimized for each individual assembly type. This is especially important for the CSNRs, which are only metastable in the current system. We envision a two-component system for future CSNRs, with one mutant to form helical rods and another mutant to form the hexagonal sheet of disks that wraps around the rod. This would minimize the formation of alternative structures, such as rod bundles, that interfere with the current system. Designing a diverse library of viral templates could have a profound impact on the synthesis of next generation functional nanostructures. Virus templating can provide a cost-effective and sustainable method to synthesize a range of advanced nanomaterials under mild conditions with nanometer precision.

## Acknowledgements

We thank Dr. Mike Strauss, Department of Anatomy and Cell Biology, McGill University, for assistance collecting electron tomography data on nanotubes.

GC and ASB were supported by the Canada Foundation for Innovation (CFI), Québec Centre for Advanced Materials (QCAM), and Natural Sciences and Engineering Research Council (NSERC). ASB also acknowledges support from the research Foundation Flanders (FWO) for a research stay at KU Leuven.

GM and SDF acknowledge support from KU Leuven (grant C1 C14/23/090) and the research Foundation Flanders - FWO (grant I006922N).

We gratefully acknowledge Polish high-performance computing infrastructure PLGrid (HPC Center: ACK Cyfronet AGH) for providing computer facilities and support within computational grant no. PLG/2023/016577.

This publication was partially developed under the provision of the Polish Ministry and Higher Education project “Support for research and development with the use of research infra-structure of the National Synchrotron Radiation Centre SOLARIS” under contract no 1/SOL/2021/2.

