## Supplementary Information for "Extended Plasmonic Nanostructures Templated by Tobacco Mosaic Virus Coat Protein"

### Materials & Methods

#### Materials

All chemicals were reagent grade or better. 100% ethanol was purchased from Commercial Alcohols. Potassium phosphate monobasic, potassium phosphate dibasic, sodium borate, EDTA, imidazole, MES, MOPS, bis-tris, triethanolamine, and sodium chloride were purchased from Fisher Scientific. Terrific Broth and LB media were purchased from MP Biomedicals. Ampicillin and chloramphenicol were purchased from Research Products International (RPI).

#### TMVP Expression

A pET20b vector encoding the sequence of WT-TMVP was purchased from NorClone Biotech by reverting an S123C mutant plasmid kindly gifted by Prof. Matthew Francis (UC Berkeley) back to the wild-type sequence. The 6H-TMVP, 6H-2R-TMVP, and 4H-TMVP plasmids were purchased from NorClone Biotech based on the WT-TMVP plasmid. For all mutants, Tuner(DE3)pLysS competent cells (Novagen) were transformed with the vector and streaked on an LB agar plate supplemented with 100 ug/mL ampicillin and 34 ug/mL chloramphenicol. After overnight incubation at 37 $^{o}$C, a single colony was used to inoculate 10 mL of LB and grown overnight with constant shaking at 37 $^{o}$C. This saturated growth was used to make frozen glycerol stocks of transformed cells stored at -80 $^{o}$C. For a typical expression, 10 mL of LB media supplemented with 100 ug/mL ampicillin and 34 ug/mL chloramphenicol was inoculated with a small portion of cells from the frozen glycerol stock and grown overnight at 30 $^{o}$C with constant shaking at 250 RPM. A 1 mL aliquot of the resulting culture was used to inoculate 1 L of Terrific Broth, grown at 37 $^{o}$C until $OD_{600}$ = 0.6, and then grown overnight at 30 $^{o}$C. No isopropylthio-β-galactoside (IPTG) was necessary for high levels of protein expression due to a combination of leaky expression of the promoter and compromised cell growth upon induction by IPTG.[1,2] Cell pellets were harvested by centrifugation and frozen at -80 $^{o}$C.

#### WT-TMVP Purification

Frozen cell pellets were thawed on ice, resuspended in cold lysis buffer (20 mM triethanolamine, 1 mM EDTA, pH 8.0), and lysed by sonication at 50% duty cycle and 60% amplitude for 5 minutes on ice. The resulting lysate was clarified by centrifugation at 32000 RCF for 30 minutes at 4 $^{o}$C. The supernatant was collected and allowed to reach room temperature. Solid ammonium sulfate was added slowly with stirring to a final concentration of 35% (v/v). The resulting precipitate was isolated by centrifugation at 32000 RCF for 30 minutes 4 $^{o}$C and resuspended in minimal lysis buffer. The solution was then dialyzed overnight at 4 $^{o}$C against 4 L lysis buffer to remove residual ammonium sulfate. Any remaining precipitate was removed by centrifugation. The solution was diluted to a final volume of 250 mL with lysis buffer and loaded on a DEAE Sepharose anion exchange column (Cytiva) overnight at 4

°C. The protein was eluted with a 0-300 mM NaCl gradient using an AKTA Avant FPLC. Fractions showing absorbance at 280 nm were analyzed by Tris-buffered SDS-PAGE on an 8-16% gradient gel (Biorad) run in constant voltage mode at 200 V and stained with Coomassie Blue R250 (Fig. S17). Pure fractions were combined and concentrated. Pure protein was dialyzed at 4 °C against 20 mM sodium borate buffer at pH 8.5, concentrated to 2.7 mg/mL, and filter sterilized. Aliquots were frozen in LN2 and stored at -80 °C until further use.

**6H-, 4H-, & 6H-2R-TMVP Purification**

Frozen cell pellets were thawed, resuspended in cold lysis buffer (15 mM imidazole, 20 mM $KH_2PO_4$, 500 mM NaCl, pH 8.0), and lysed by sonication at 50% duty cycle and 60% amplitude for 5 minutes on ice. The resulting lysate was clarified by centrifugation at 32000 RCF for 30 minutes at 4 °C. The supernatant was loaded on a 5 mL Ni-NTA column (HisTrap HP, Cytiva). The protein was eluted with a 15-1000 mM imidazole gradient using an AKTA Avant FPLC. Fractions showing absorption at 280 nm were analyzed by Tris-buffered SDS-PAGE on an 8-16% gradient gel run in constant voltage mode at 200 V and stained with Coomassie Blue R250 (Fig. S17). Pure fractions were combined and concentrated. Pure protein was dialyzed at 4 °C against 20 mM sodium borate buffer with 50 mM EDTA at pH 9.0, concentrated to 2.8 mg/mL, and filter sterilized. Aliquots were frozen in LN2 and stored at -80 °C until further use.

**3 nm AuNP Synthesis**

3 nm AuNPs were synthesized as previously reported.[3] 19.78 mL of Milli-Q water, 100 µL of 50 mM sodium citrate, and 93.27 µL of 53.61 mM $HAuCl_4$ were combined with vigorous stirring in a 25 mL Erlenmeyer flask at room temperature. 600 µL of freshly prepared cold 100 mM sodium borohydride solution was rapidly injected. The solution was stirred vigorously for 3 hours after which 10 mg of BSPP was added as the capping agent. The mixture was allowed to stir overnight. The AuNPs were washed by repeatedly concentrating and diluting with Milli-Q water in spin concentrator tubes. The final dilution was performed with a solution of 10 mg of BSPP in Milli-Q water to obtain an AuNP concentration of approximately 2.5 µM. For size distributions see Fig. S15A.

**5 nm AuNP Synthesis**

5 nm AuNP synthesis was based on a previously reported method.[4] 100 µL of 53.6 mM $HAuCl_4$ was combined with 9.25 mL of Milli-Q water with vigorous stirring in a 20 mL vial at room temperature. 650 µL of freshly prepared 53.6 mM $NaBH_4$ solution was rapidly injected. After 1 minute, the vial was placed in boiling water for 3 minutes and then allowed to cool to room temperature. The AuNPs were mixed with 10 mg BSPP and stirred overnight. The particles were then washed and concentrated to 1.5 µM just as with the 3 nm AuNPs. For size distributions see Fig. S15B.

**8 nm AuNP Synthesis**

Particles were synthesized as previously reported.[3] 3 nm seed particles were first synthesized by combining 46.4 mL of Milli-Q water with 3.2 mL of 34.3 mM sodium citrate, 33.3 mL of 18 mM tannic acid, and 333.0 mL of 150 mM sodium carbonate in a 100 mM Erlenmeyer flask with vigorous stirring at 70 °C. 155.45 mL of 53.6 mM tetrachloroauric acid was added and stirred vigorously at 70 °C for 30 minutes. The solution was cooled, and the volume was adjusted to 50 mL. 4 mL of 200 mM Tris base was added to 176 mL of Milli-Q water and stirred vigorously at 70 °C. 186.55 mL of 53.6 mM tetrachloroauric acid and 3 nm particle seed solution (20 mL) were quickly added and stirred vigorously at 70 °C for 30 min or until the color stabilized. The solution was removed from heat and stirred for an additional 2–3 h. BSPP was added to a final concentration of 1 mg/mL, and the solution was stirred overnight. Filtered saturated NaCl was added dropwise until the solution turned dark gray, indicating precipitation of the AuNPs. The precipitated particles were pelleted in an Eppendorf MiniSpin centrifuge, and the supernatant was discarded. Nanoparticles were resuspended to a concentration of 1.5 uM in an aqueous solution of 1 mg/mL BSPP.

**Protein Self-Assembly and AuNP Attachment**

All samples were prepared by dialysis of stock protein diluted to the desired concentration. Additives, such as imidazole or ethanol, were included in the protein solution before dialysis to prevent large volume changes from osmotic pressure and to ensure that the additives affected the full duration of protein assembly.

**Hexagonal Sheets**

Hexagonally packed sheets of 6H-TMVP disks were formed by overnight dialysis of the protein at 2.5 mg/mL from pH 9.0 to pH 6.0 in 100 mM potassium phosphate, 5.0 mol% ethanol at room temperature. Samples were adjusted to the correct ethanol concentration before dialysis. Dialyzed samples were incubated at room temperature for 3 days to allow large sheets to form. Alternatively, protein at 1.5 mg/mL was dialyzed against 50 mM potassium phosphate, 500 mM imidazole, pH 6.8 and incubated for at least 2 days. In either case, single layer sheets were stable without multilayer aggregation for at least 1 week. AuNPs were attached to hexagonally packed sheets by addition of 60 µL of 3 or 5 nm AuNPs at stock concentrations to 10 µL of assembled 6H-TMVP. AuNP attachment was better in the ethanol samples than the imidazole samples. The mixtures were thoroughly mixed by pipette and left to assemble overnight at room temperature. Excess AuNPs could be removed by allowing large structures to settle out of solution and then carefully replacing the supernatant with an equal volume of fresh buffer.

**Square Sheets**

6H-TMVP at 2.5 mg/mL was dialyzed overnight to pH 6.0 in 100 mM potassium phosphate at 4 ºC. To initiate sheet formation, 10 µL of 6H-TMVP was mixed with 60 µL of 3 or 5 nm AuNPs at 4 ºC. Samples were incubated at 4 ºC for 2 days. While unbound AuNPs could be removed as described above, this was found to also reduce the number of particles in each ring.

**Imidazole-induced Fibers**

6H-TMVP at 1.5 mg/mL was dialyzed overnight against pH 6.8, 50 mM potassium phosphate, 100 mM imidazole at 4 ºC. Samples were incubated at 4 ºC for 2 days to allow fiber formation. Fibers were also formed at higher protein concentrations by keeping the same protein to imidazole ratio.

**Core-Shell Nanorods**

100 µL of 6H-TMVP at 1.0 mg/mL was dialyzed overnight against 50 mM potassium phosphate, 3.5 mol% ethanol, pH 6.0 at 4 °C. Samples were then incubated at room temperature for at least 3 days to form core-shell rods. Samples failed to form CSNRs when left at 4 °C for one week without any time at room temperature. Incubation at room temperature for less than 3 days typically resulted in rods surrounded by loosely associated disks lacking a well-ordered sheet structure.

### Characterization

#### Transmission Electron Microscopy

Room temperature transmission electron microscopy was performed on a 200 kV ThermoScientific Talos F200X at the Facility for Electron Microscopy Research (FEMR) at McGill University. All samples were deposited on either plain carbon or ultrathin carbon on lacey carbon support grids. Samples were typically deposited for 60 seconds, wicked and washed with three drops of ultrapure water. Samples containing large sheets were deposited for 3 minutes to compensate for much lower effective concentration after self-assembly. Protein was stained with 3% uranyl acetate or 2% phosphotungstic acid for 60 seconds after washing. Nanoparticle-containing samples were not stained unless otherwise noted. Electron tomography was carried out on a negatively stained sample with a single axis tilt from +60 to -60°. Tomographic reconstruction was performed with Etomo in the IMOD package.[5]

#### Optical Measurements

Darkfield hyperspectral images were collected using an inverted LIMA imaging system (Photon Etc.) equipped with a supercontinuum FIU-15 Laser (NKT Photonics.) and LLTF Contrast tunable filter (Photon Etc). The image was collected through an Eclipse Ti-2U microscope (Nikon) equipped with a Dark Field Condenser 0.95 – 0.80 and 60X S Plan Fluor Objective (Nikon) onto an ORCA-Flash4.0 Digital CMOS camera (Hamamatsu). The spectra were corrected against the dark counts of the imaged slide, followed by normalization with the power profile signature of the laser taken prior to sample collection.

Correlations were accomplished using pattern matching, with grid co-localization accomplished using pinpointer grids.

#### Atomic Force Microscopy

A sample of assembled 6H-TMVP was diluted with buffer by a factor of ten and a volume of 10 µL was deposited onto highly oriented pyrolytic graphite (HOPG). Subsequently, 10 µL of water (high purity ICP-MS grade) was added. After 10 minutes, the droplet was removed using tissue, and the sample was dried under argon.

AFM imaging was performed using a Multimode 8 (Bruker) with a Nanoscope V controller in tapping mode under ambient conditions. OMCL-AC240TS-R3 probes (spring constant ≈ 2 N/m) with a resonance frequency of approximately 70 kHz were employed. Gwyddion software was used for AFM image processing.

#### Cryogenic Electron Microscopy

4 uL of a solution of 6H-TMVP at 2.0 mg/mL in 50 mM potassium phosphate pH 6.0 was applied to a glow discharged grid and plunge frozen in liquid ethane using an FEI Vitrobot. 8488 micrographs were collected using a Titan Krios cryo-microscope with a Falcon III camera at 75k magnification. All micrographs were motion-corrected using MotionCorr2 and CTF estimation was performed using PatchCTF in cryoSPARC v4.3.0.[6] Automated particle picking was performed using the BlobPicker tool from 500 micrographs. Approximately 3400 particles were extracted and classified into 2D classes. These classes were then used for another round of particle picking and 2D classification using template picker from 500 micrographs. The best classes were used as a template for template picker on the full set of micrographs. 2D classes from the full data set were used to train a Topaz neural network. Two rounds of training were performed. Each training was followed by particle extraction, and 2D classifications to select the best-looking classes. *Ab-initio* reconstruction was performed on 102,722 particles from 5 classes. The best 4 classes were subjected to homogeneous refinement with C1 symmetry followed by another round of homogeneous refinement with $C_{17}$ symmetry. Final maps had resolutions from 3-4 Å based on FSC at 0.143 threshold.

### Figures

#### Imidazole Induced Bundles

Bundles formed in the presence of imidazole have visible striations, although it is unclear if the striations are directly caused by stacked disks or an interference pattern from overlapping stacked disks/helical rods. Bundles reaching up to 7 µm in length with an aspect ratio of 23 were observed. Bundles were formed at a wide range of protein concentrations by maintaining a similar imidazole to protein ratio. It is unclear what role imidazole plays in the formation of these fibers. Bundles were not observed under the same solution conditions without imidazole, or at high ionic strength from NaCl. Similar bundles were formed by a mutant Trp RNA binding protein (TRAP), which self-assembles into rings somewhat similar to TMVP disks.[7]

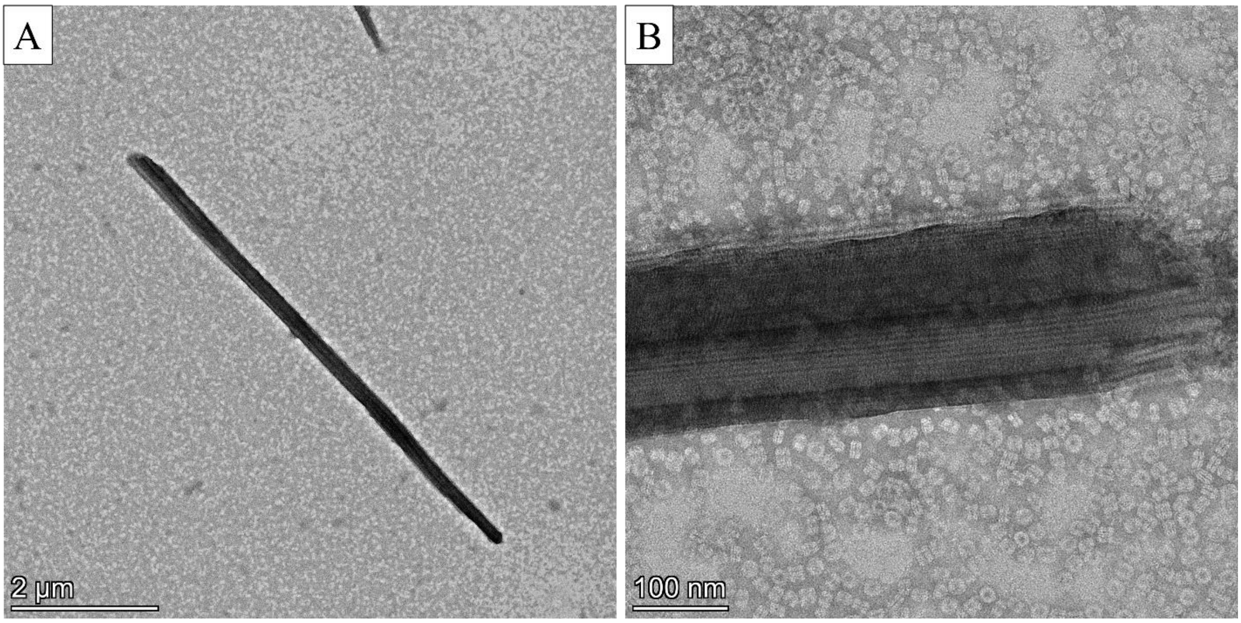


**Figure S1.** TEM images of long bundles of rods/stacked disks formed in the presence of imidazole.

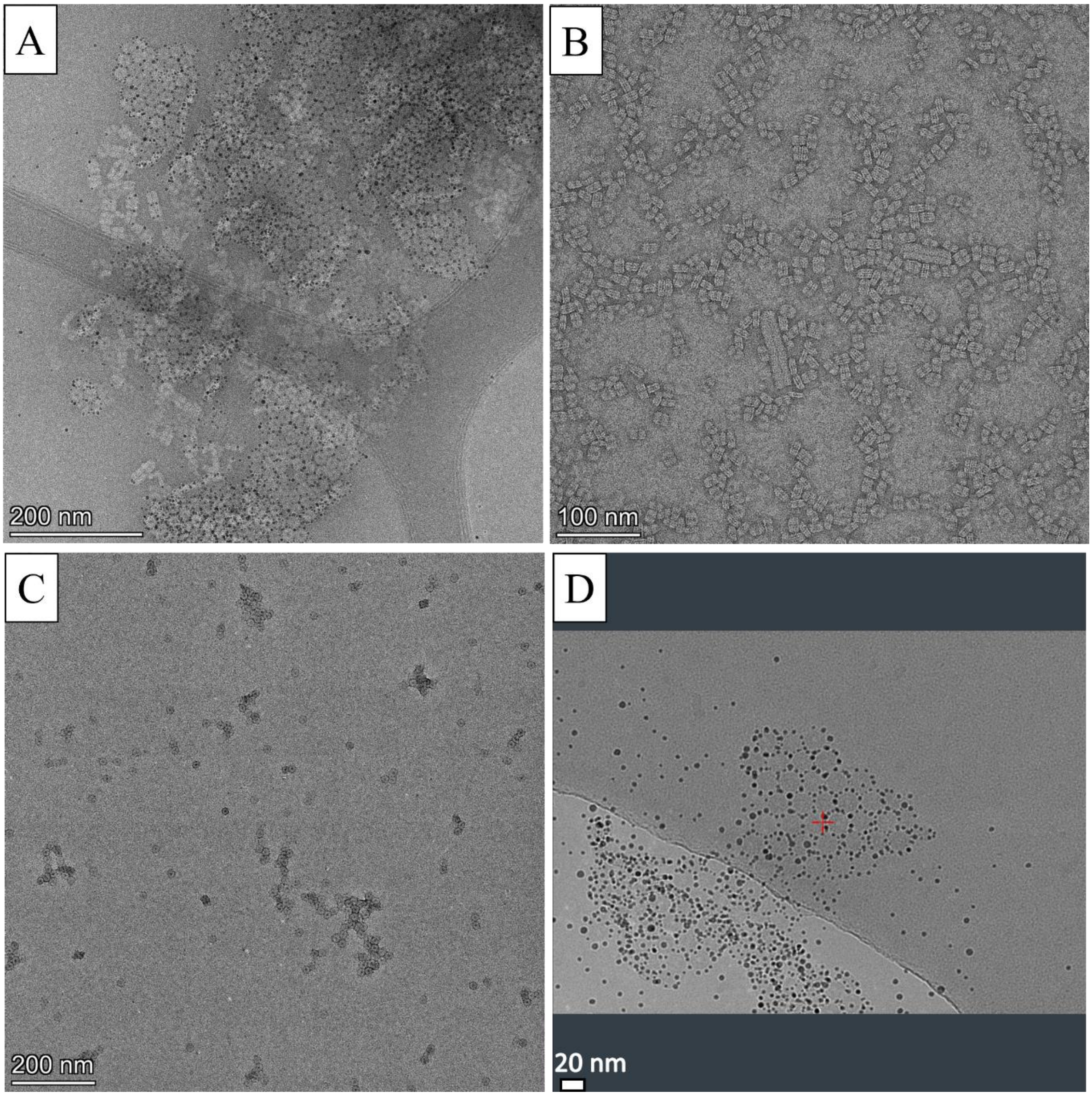


**Figure S2.** (A) TEM image of sheets under square conditions assembled directly on a TEM grid with negative staining. (B) Protein under square sheet conditions before addition of AuNPs. (C) Protein under square conditions after addition of only BSPP solution. (D) Hexagonal sheet with AuNPs attached in cryo-EM.

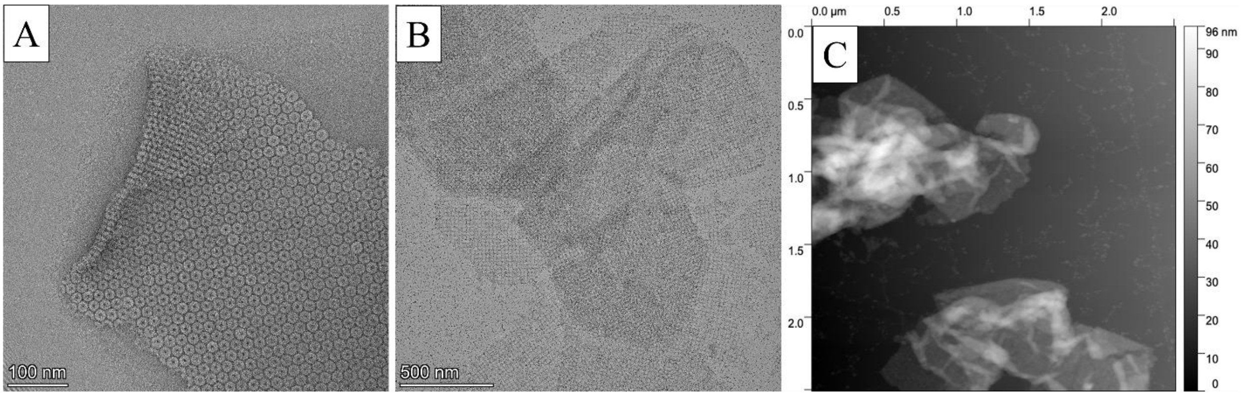


**Figure S3.** Images of folded sheets. (A) TEM image of folded hexagonal sheet. (B) TEM image of exfoliated/folded square sheets. (C) AFM image of folded sheets.

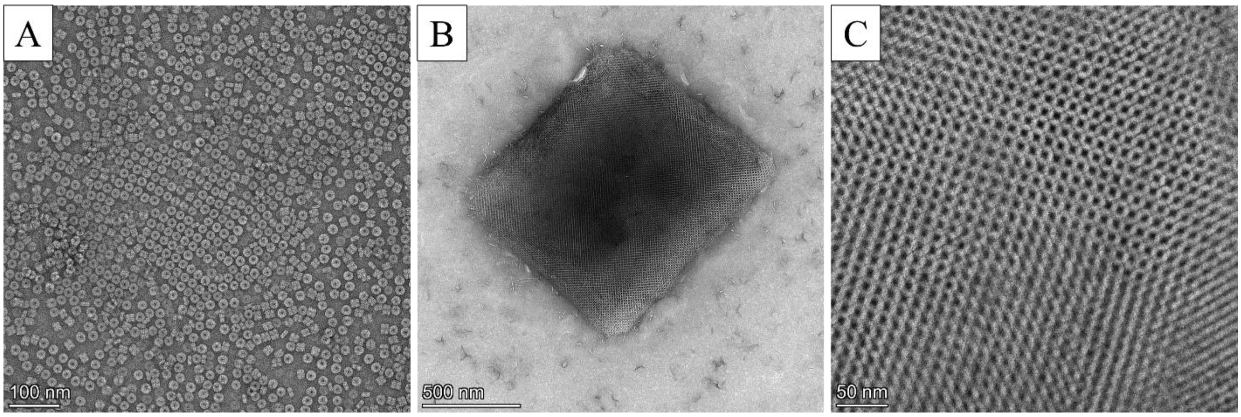


**Figure S4.** TEM images of 6H-2R-TMVP square sheets. (A) Single-layer square sheets spontaneously form on TEM grids. (B) Crystal with apparent body-centered tetragonal packing. (C) High mag image of BCT crystal.

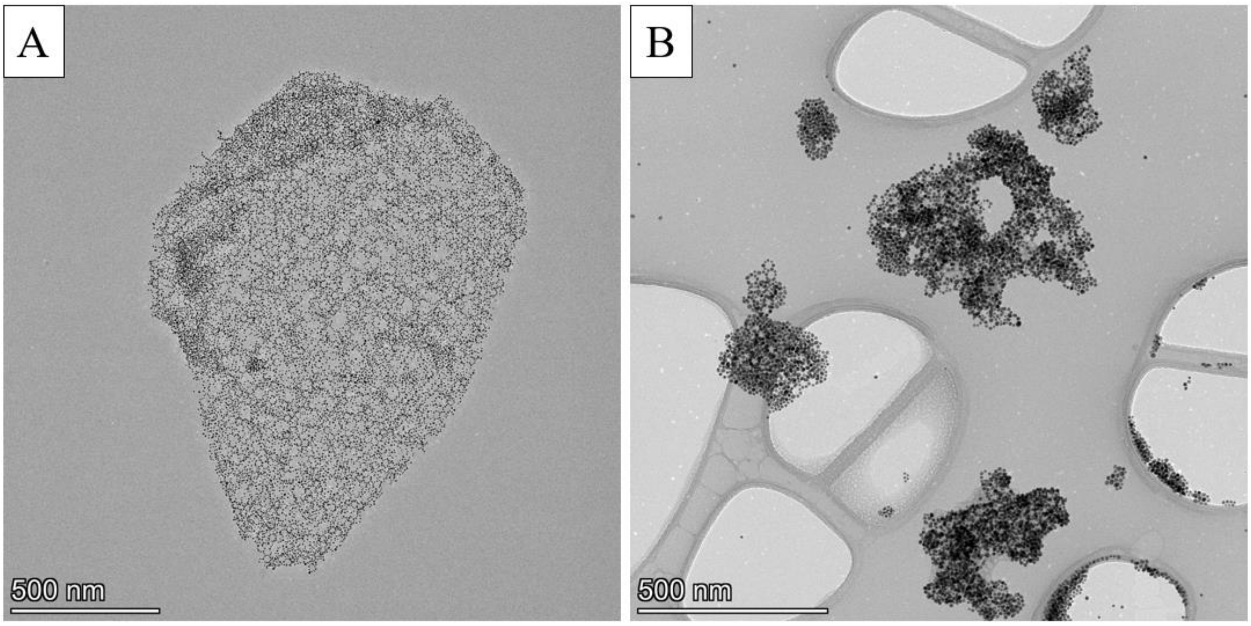


**Figure S5.** TEM images of (A) Hexagonal sheet with 5 nm AuNPs. (B) Failed hexagonal sheets formed with 8 nm AuNPs.

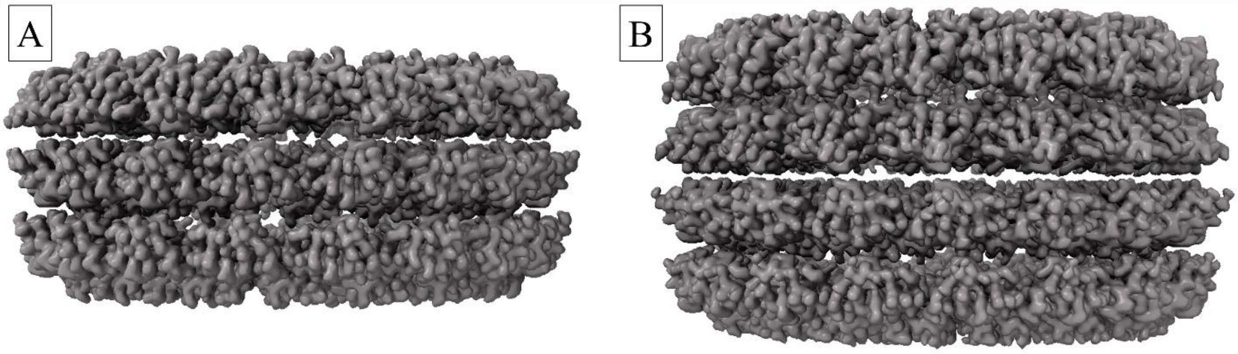


**Figure S6.** Side view of cryo-EM maps of 3- and 4-layer 6H-TMVP disks created with UCSF ChimeraX.[8]

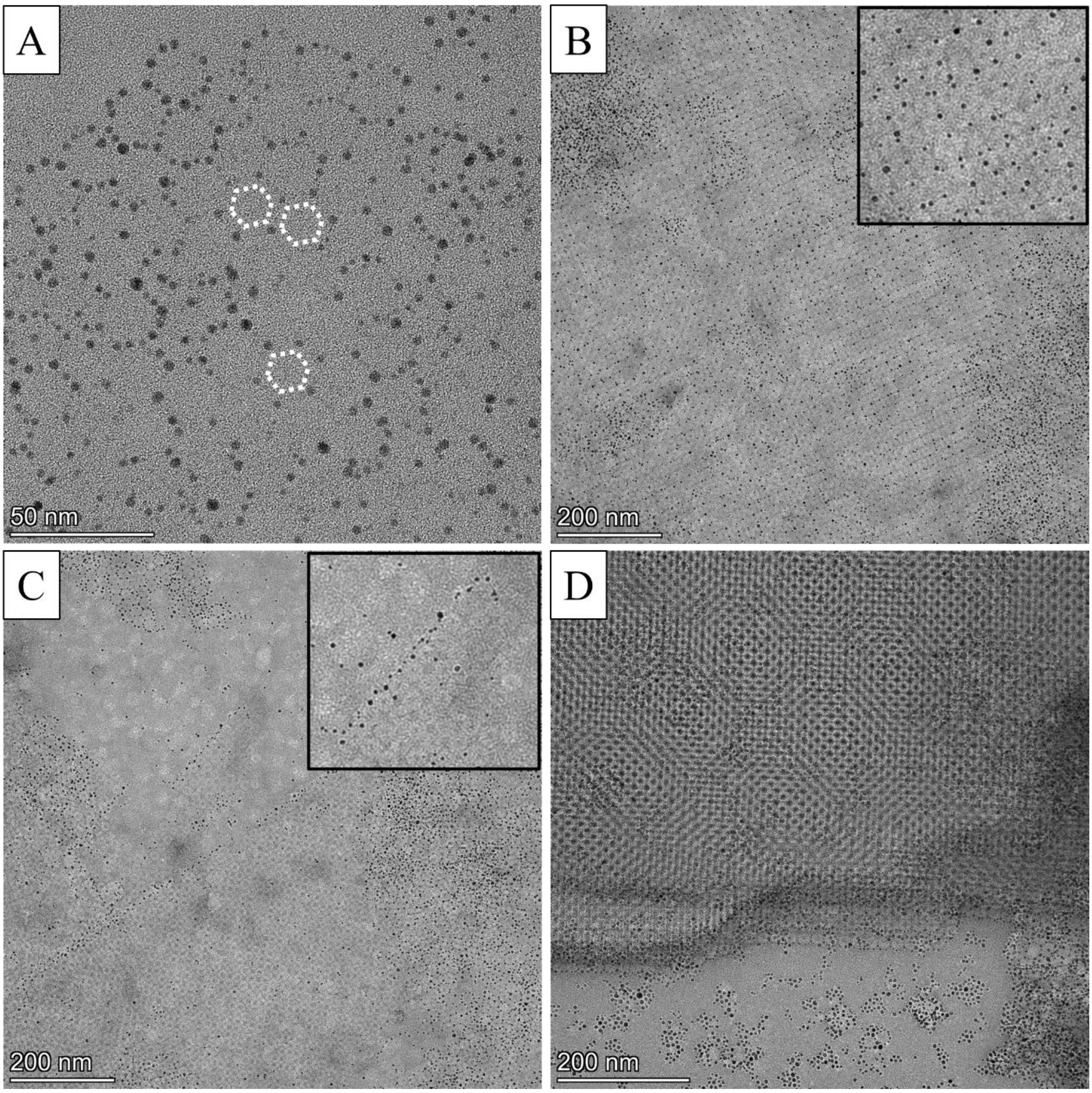


**Figure S7.** TEM images of (A) 6H-TMVP hexagonal sheets 1 hour after mixing with AuNPs, (B-D) Negatively stained square sheet sample 15 minutes after addition of 3 nm AuNPs showing (B) AuNPs binding at the vertices of the square lattice, (C) AuNPs binding to the outer edge of square sheets, and (D) multilayer sheets that appear to be mostly lacking bound AuNPs.

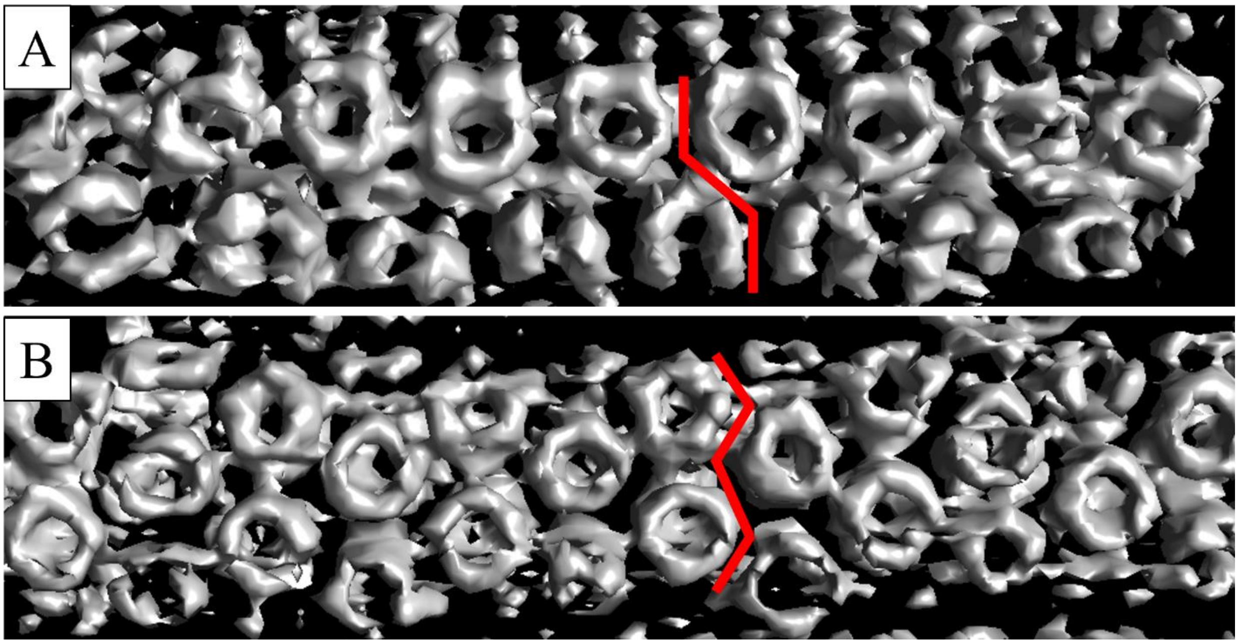


**Figure S8.** Negative stain electron tomograms showing (A) armchair and (B) zigzag 6H-TMVP nanotubes.

### EDTA/High Salt Controls

Although protein was treated with ethylenediaminetetraacetic acid (EDTA) after purification, assembly was also attempted in the presence of EDTA to test the possibility that self-assembly was due to metal contamination. CSNRs successfully formed with 50 mM EDTA in the assembly buffer (Fig. S9). Protein assembled under hexagonal sheet conditions with 50 mM EDTA did not form hexagonal sheets. Instead, the disks mainly formed small, disordered aggregates and a small population of crystalline particles which appeared to be monoclinic. In 200 mM EDTA and hexagonal sheet conditions, only small, disordered aggregates were observed. TMVP assembled under square sheet conditions with 50 mM EDTA did not show any notable differences compared to the protein without EDTA. Square sheet conditions with 200 mM EDTA caused the disks to form disordered aggregates as well as polycrystalline assemblies with packing that appeared to be either monoclinic or body-centered tetragonal depending on the region (Fig. S9).

Addition of AuNPs to the protein in hexagonal conditions with 50 mM EDTA led to the formation of a mixture of small, disordered aggregates and square sheets (Fig. S10). The square sheets obtained by this method were much smaller and lower quality than those from standard square sheet conditions. The conditions for hexagonal and square sheets are quite similar, so this result is not entirely surprising. When mixed with AuNPs, the protein under hexagonal conditions with 200 mM EDTA assembled into poorly formed hexagonal sheets and random aggregates. Protein in square sheet condition with 50 or 200 mM EDTA formed low quality hexagonal sheets upon addition of AuNPs. The sheets were larger with more complete AuNP binding in 50 mM EDTA compared to 200 mM EDTA. The structures formed in high EDTA concentrations are similar to those formed under high salt concentrations, suggesting that the effects may be due to ionic strength rather than metal chelation (Fig. S11).

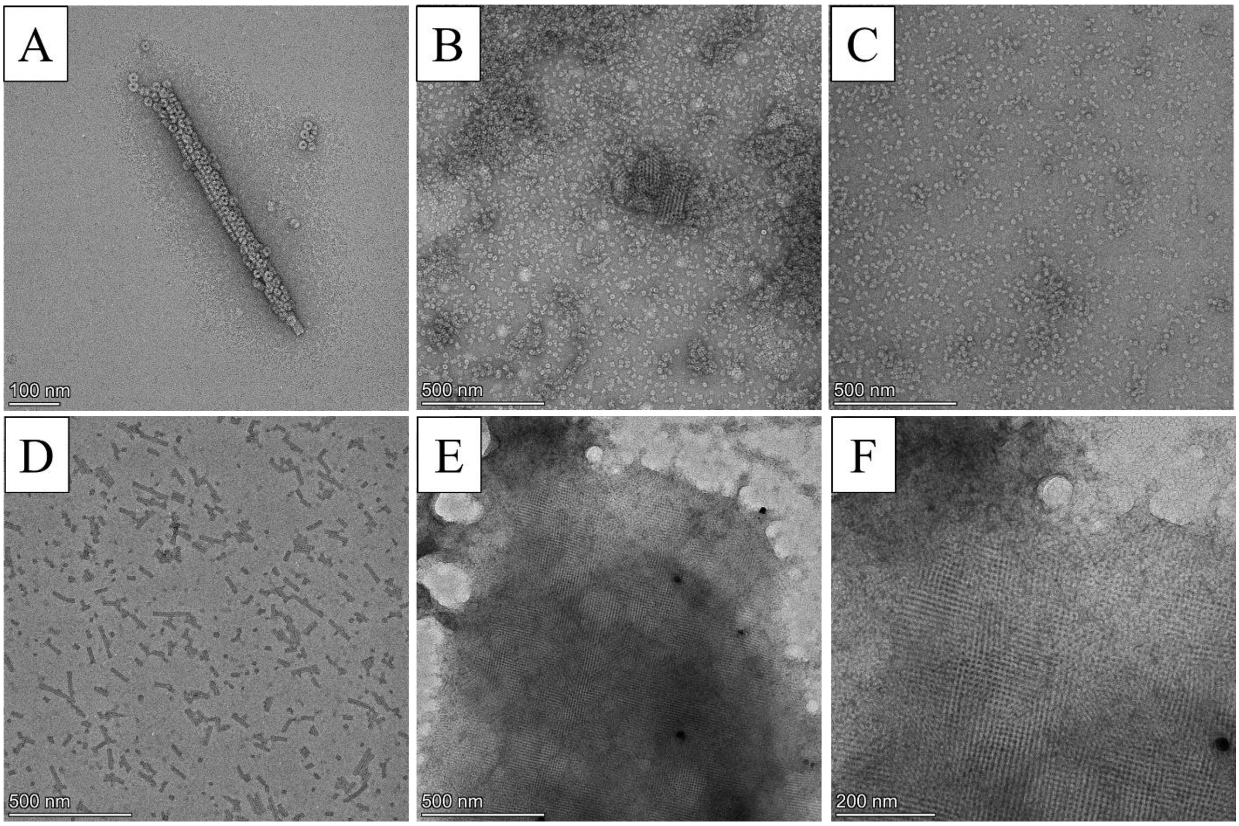


**Figure S9.** TEM images of (A) CSNR formed in 50 mM EDTA. (B) 6H-TMVP in hexagonal sheet conditions in 50 mM EDTA. (C) 6H-TMVP in hexagonal sheet conditions s in 200 mM EDTA. (D) 6H-TMVP in square sheet conditions in 50 mM EDTA. (E-F) 6H-TMVP in square sheet conditions in s 200 mM EDTA.

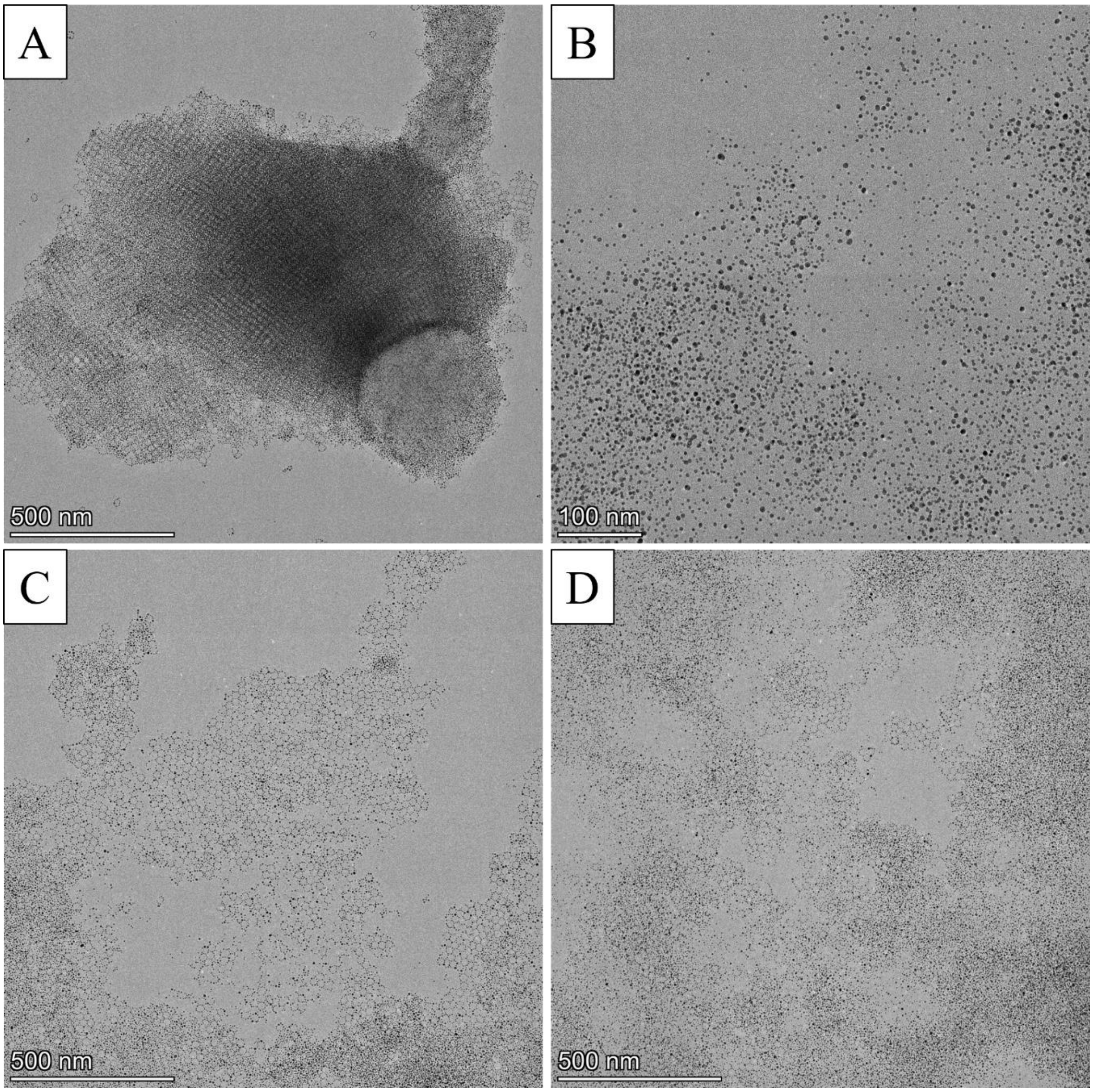


**Figure S10.** TEM images of EDTA controls after addition of AuNPs. (A) 50 mM EDTA hexagonal conditions. (B) 200 mM EDTA hexagonal conditions. (C) 50 mM EDTA square conditions. (D) 200 mM EDTA square conditions.

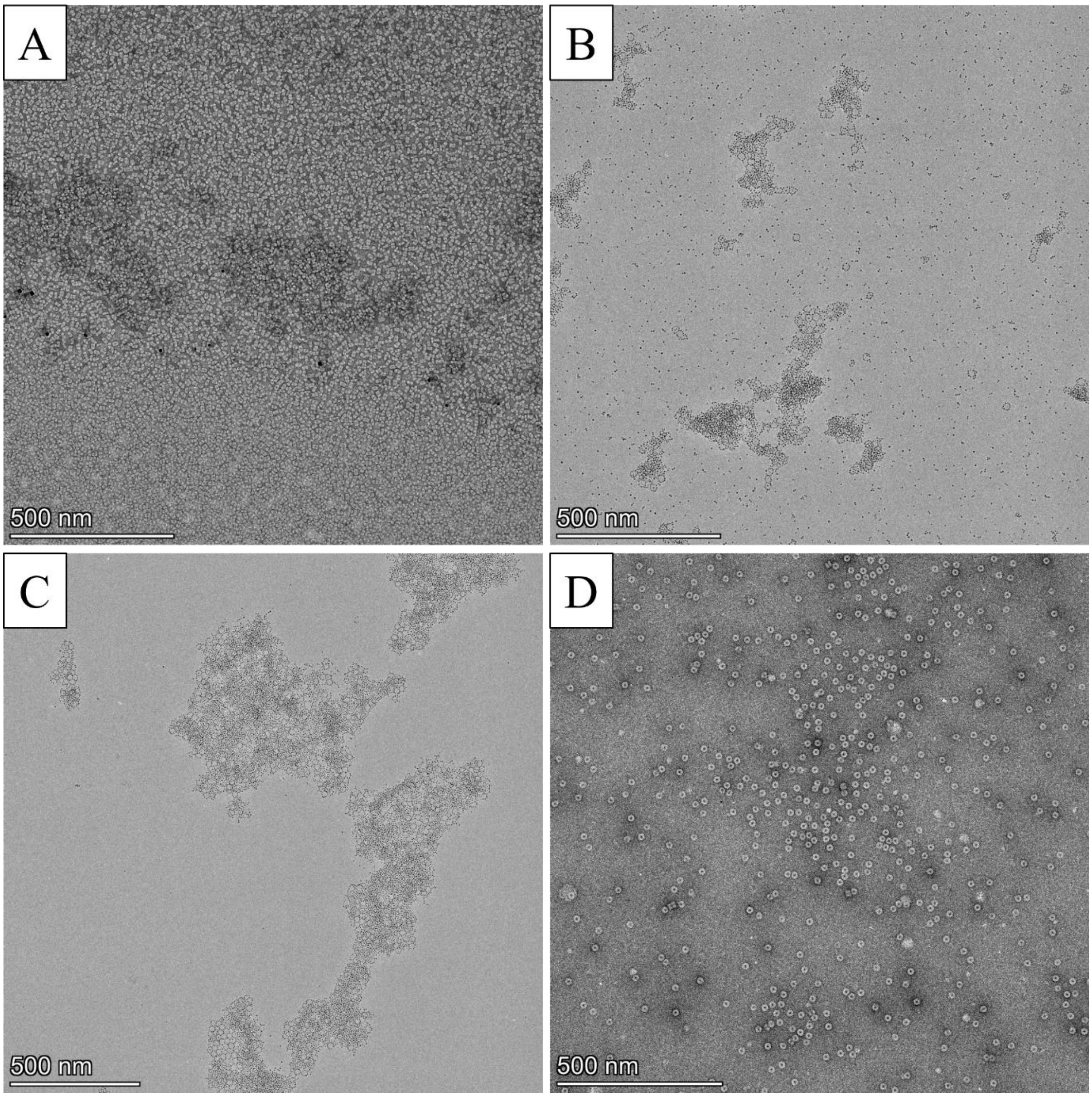


**Figure S11.** TEM images of (A) 500 mM NaCl hexagonal conditions. (B) 500 mM NaCl hexagonal conditions with AuNPs. (C) 500 mM NaCl square conditions with AuNPs. (D) 4H-TMVP does not assemble without metal ions.

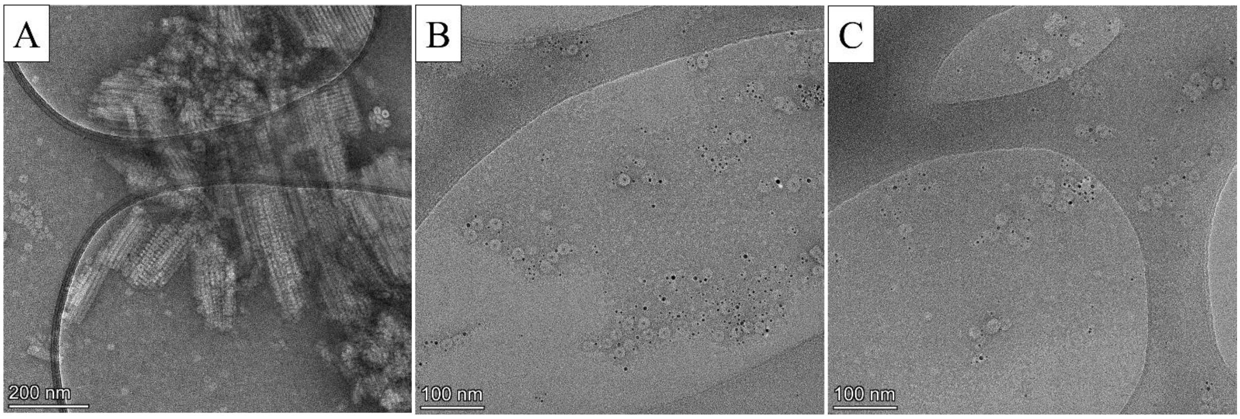


**Figure S12.** TEM images of (A) 6H-TMVP assembled in 70 mM sodium sulfate, 20 mM bis-tris, pH 6.0. (B) WT-TMVP mixed with AuNPs under hexagonal sheet conditions. (C) WT-TMVP mixed with AuNPs under square sheet conditions. No AuNP rings were observed in WT-TMVP samples.

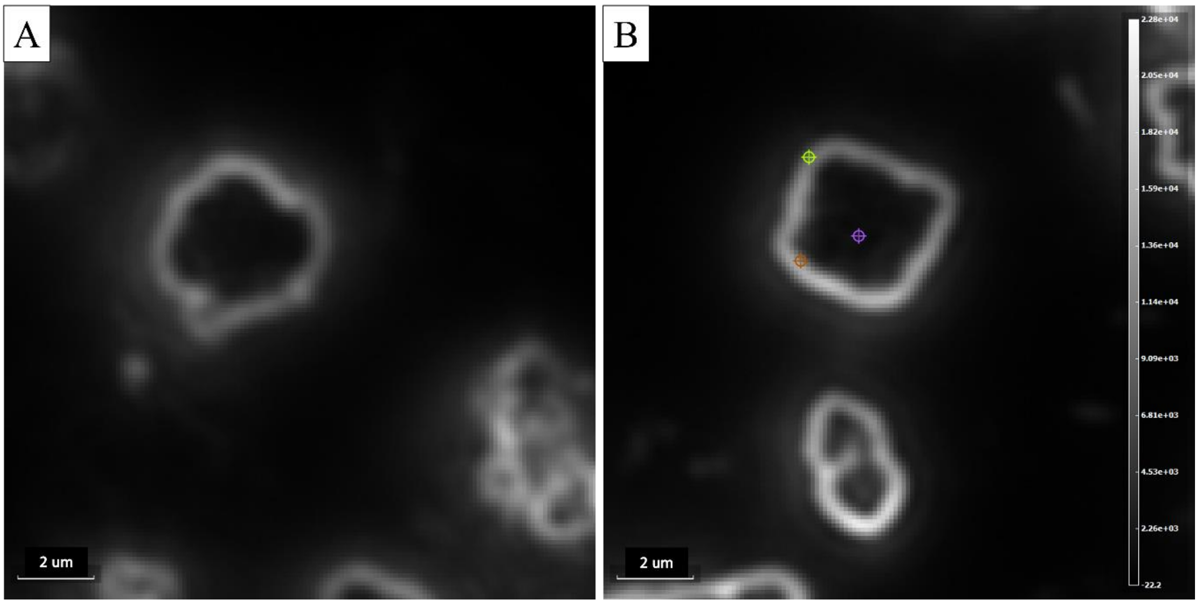


**Figure S13.** Hyperspectral darkfield microscope image of (A) hexagonal sheets and (B) square sheets showing bright edges and dark interior. Coloured targets are from spectra selection.

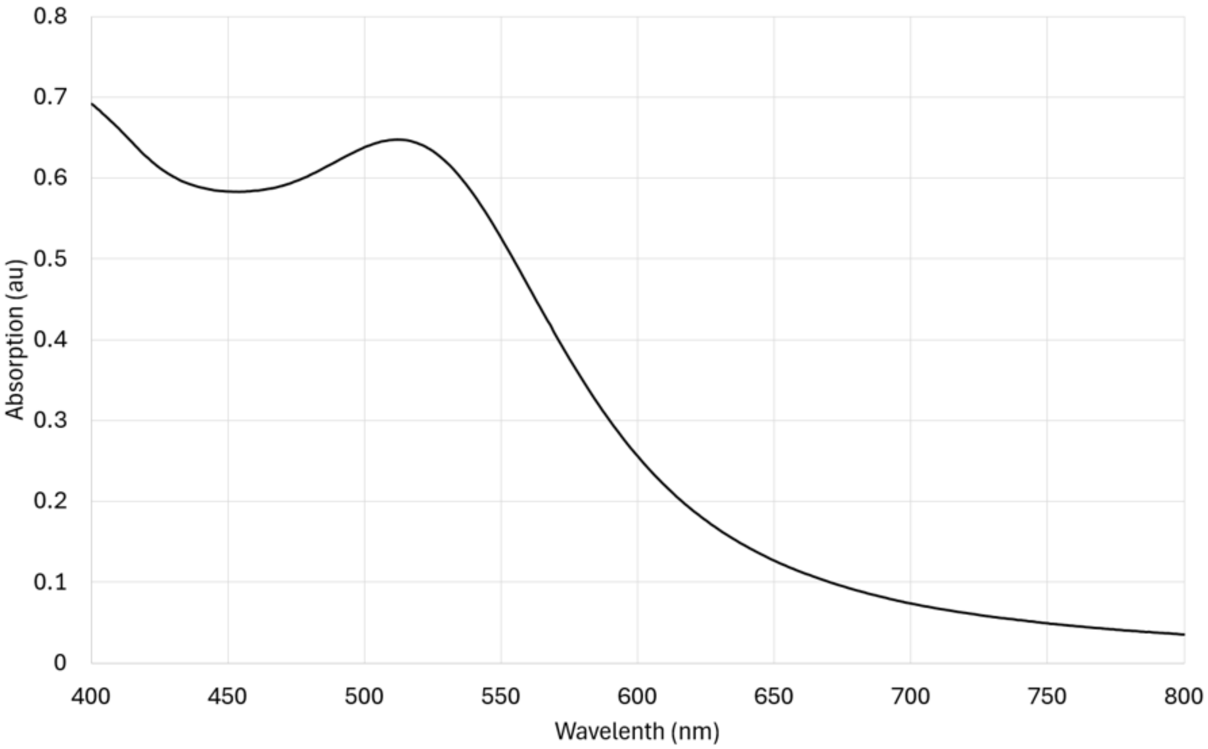


**Figure S14.** Solution UV-vis spectrum of 3 nm AuNPs showing absorption peak at 514 nm.

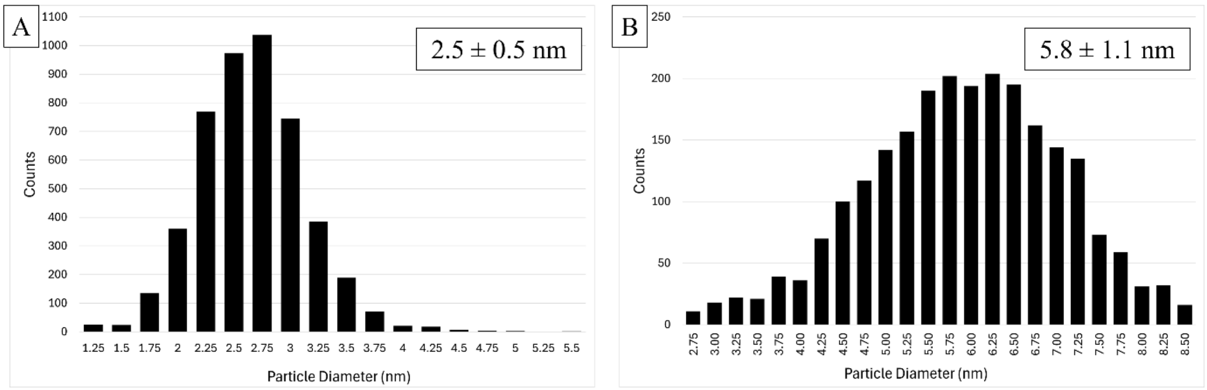


**Figure S15.** Particle size distributions of (A) 3 nm AuNPs, (B) 5 nm AuNPs.

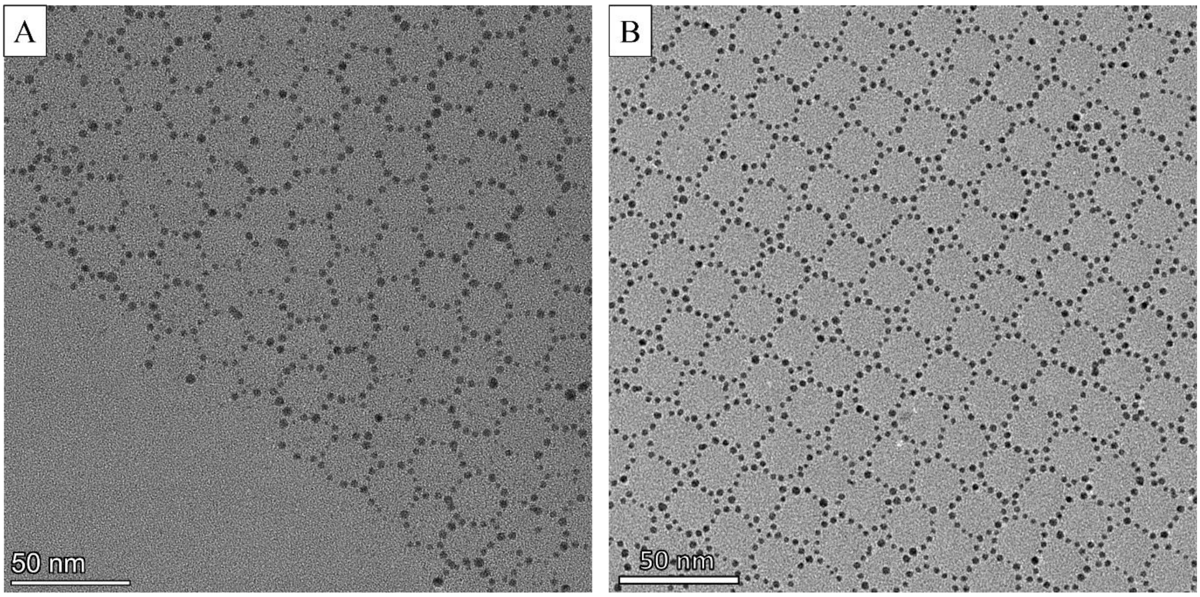


**Figure S16.** High magnification images of (A) hexagonal and (B) square sheets.

| Measurement | Average | SD |
|---|---|---|
| Hexagonal interparticle distance | 4.30 | 0.92 |
| Square interparticle distance | 4.53 | 0.82 |
| Hexagonal ring center-to-center | 20.45 | 1.74 |
| Square ring center-to-center | 20.67 | 1.50 |

| Rings without central particle | Percent |
|---|---|
| Hexagonal | 77.0 |
| Square | 80.6 |

**Table S1.** Measurement and statistics of spacing between adjacent AuNPs in rings, center-to-center distances between disks in sheets, and the presence of AuNPs in the centers of the nanorings. SD is standard deviation.

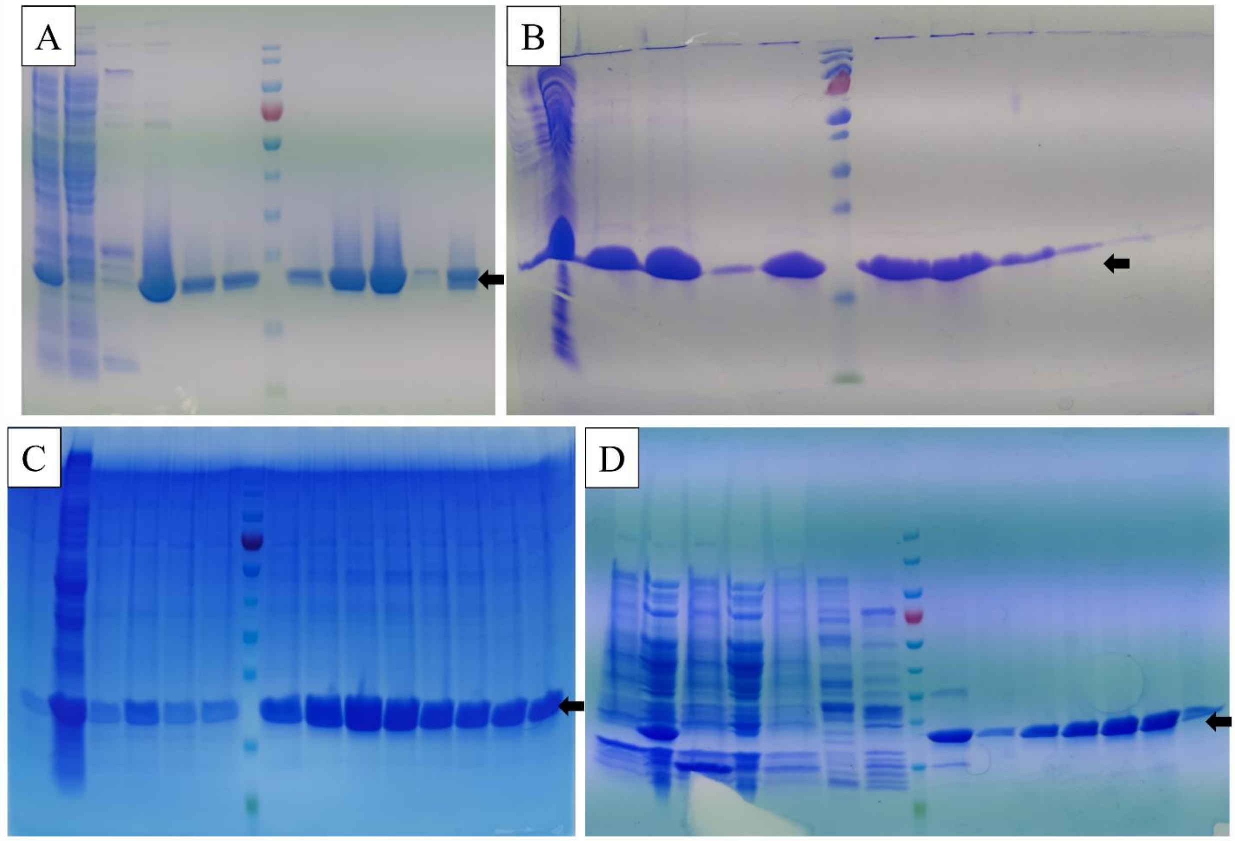


**Figure S17.** SDS-PAGE of cell lysates and column elutions of (A) 6H-TMVP, (B) 4H-TMVP, (C) WT-TMVP, (D) 6H-2R-TMVP. Molecular weight markers: 10 (green), 15, 25, 35, 40, 55, 70 (red), 100, 130, 180 kDa. Arrows indicate the mass of the target protein.
